# Inhibition of the αv integrin-TGF-β axis improves natural killer cell function against glioblastoma stem cells

**DOI:** 10.1101/2020.03.30.016667

**Authors:** Hila Shaim, Mayra Hernandez Sanabria, Rafet Basar, Fang Wang, May Daher, Daniel Zamler, Joy Gumin, Konrad Gabrusiewicz, Qi Miao, Jinzhuang Dou, Abdullah Alsuliman, Lucila Nassif Kerbauy, Sunil Acharya, Vakul Mohanty, Pinaki Banerjee, Mayela Mendt, Sufang Li, Junjun Lu, Jun Wei, Natalie Wall Fowlkes, Elif Gokdemir, Emily L. Ensley, Mecit Kaplan, Nadima Uprety, Cynthia Kassab, Li Li, Gonca Ozcan, Yifei Shen, April L. Gilbert, Mustafa Bdiwi, Ana Karen Nunez Cortes, Enli Liu, Jun Yu, Nobuhiko Imahashi, Luis Muniz-Feliciano, Jian Hu, Giulio Draetta, David Marin, Dihua Yu, Stephan Mielke, Matthias Eyrich, Richard Champlin, Ken Chen, Frederick F. Lang, Elizabeth Shpall, Amy Heimberger, Katayoun Rezvani

## Abstract

Glioblastoma, the most aggressive brain cancer, often recurs because glioblastoma stem cells (GSCs) are resistant to all standard therapies. Here, we show that patient-derived GSCs, but not normal astrocytes, are highly sensitive to lysis by healthy allogeneic natural killer (NK) cells *in vitro*. In contrast, single cell analysis of autologous, tissue infiltrating NK cells isolated from surgical samples of high-grade glioblastoma patient tumors using mass cytometry and single cell RNA sequencing revealed an abnormal phenotype associated with impaired lytic function compared with peripheral blood NK cells from GBM patients or healthy donors. This immunosuppression was attributed to an integrin-TGF-β mechanism, activated by direct cell-cell contact between GSCs and NK cells. Treatment of GSC-engrafted mice with allogeneic NK cells in combination with inhibitors of integrin or TGF-β signaling, or with *TGF-β receptor 2* gene-edited NK cells prevented GSC-induced NK cell dysfunction and tumor growth. Collectively, our findings reveal a novel mechanism of NK cell immune evasion by GSCs and implicate the integrin-TGF-β axis as a useful therapeutic target to eliminate GSCs in this devastating tumor.

## INTRODUCTION

Glioblastoma multiforme (GBM) or grade IV astrocytoma, is the most common and aggressive type of primary brain tumor in adults. Despite current treatment with resection, radiotherapy and temozolamide, the outcome is poor with a reported median survival of 14.6 months and a 2-year survival of 26.5% as the tumor invariably relapses^1, 2^. This dismal outcome has stimulated keen interest in immunotherapy as a means to circumvent one or more of the factors that have limited the impact of available treatments: (i) rapid growth rate of these aggressive tumors; (ii) their molecular heterogeneity and propensity to invade critical brain structures, and (iii) the tumor regenerative power of a small subset of glioblastoma stem cells (GSCs)^3, 4^.

Emerging results from preclinical studies support the concept that GBM tumors and their associated stem cells may be susceptible to immune attack by natural killer (NK) cells^5, 6,7,8,9^. These innate lymphocytes have a broad role in protecting against tumor initiation and metastasis in many types of cancer, and they have distinct advantages over T cells as candidates for therapeutic manipulation^10, 11^. However, the vast majority of tumor cells that have been studied to date possess defenses, allowing them to evade NK cell-mediated cytotoxicity. These include disruption of receptor-ligand interactions between NK and tumor cells and the release of immunosuppressive cytokines into the microenvironment, such as TGF-β^12, 13, 14, 15^. Even if one could shield NK cells from the evasive tactics of GBM tumors, it may not be possible to eradicate a sufficient number of self-renewing GSCs to sustain complete responses. Indeed, very little is known about the susceptibility of GSCs to NK cell surveillance *in vivo*. Thus, to determine if GSCs can be targeted by NK cells *in vivo*, we designed a preclinical study and used single cell analysis of primary GBM tissue from patients undergoing surgery to determine the extent to which NK cells infiltrate sites of active tumor and the potency with which they eliminate patient-derived GSCs.

Here, we have shown that NK cells comprise one of the most abundant lymphoid subsets infiltrating GBM tumor specimens but possess an altered NK cell phenotype that correlates with reduced cytolytic function, indicating that GBM tumors generate a suppressive microenvironment to escape NK cell antitumor activity. GSCs proved highly susceptible to NK-mediated killing *in vitro*, but evaded NK cell recognition via a mechanism requiring direct αv integrin-mediated cell-cell contact, leading to the release and activation of TGF-β by the GCSs. In a patient-derived xenograft (PDX) mouse model of glioblastoma, GSC-induced NK dysfunction was completely prevented by integrin or TGF-β blockade or by CRISPR gene editing of the TGF-β receptor 2 (*TGFβR2)* on NK cells, resulting in effective control of the tumor. Taken together, these data suggest that inhibition of the αv integrin-TGF-β axis could overcome a major obstacle to effective NK cell immunotherapy for GBM.

## RESULTS

### GSCs are susceptible to NK cell-mediated killing

The GSCs can be distinguished from their mature tumor progeny at the transcriptional, epigenetic and metabolic levels^16, 17^, raising the question of whether these cells can be recognized and killed by NK cells. We therefore asked whether patient-derived GSCs, defined as being capable of self-renewal, pluripotent differentiation, and tumorigenicity when implanted into an animal host, are susceptible to NK cell cytotoxic activity as compared with healthy human astrocytes. We derived GSCs from patients with various glioblastoma subtypes including mesenchymal (GSC20, GSC267), classical (GSC231, GSC6-27), and proneural (GSC17, GSC8-11, GSC262) while also showing heterogeneity in the O(6)-Methylguanine-DNA methyltransferase (MGMT) methylation status (methylated: GSC231, GSC8-11, GSC267; indeterminate: GSC6-26, GSC17, GSC262). K562 targets were used as positive control because of their marked sensitivity to NK cell mediated killing due to lack of expression of HLA class I^18^. Across all effector:target (E:T) ratios, healthy donor NK cells killed GSCs (n=6) and K562 cells with equal efficiency and much more readily than healthy human astrocytes (n=6), which displayed a relative resistance to NK cell-mediated killing **(Fig. 1A)**. Multi-parametric flow cytometry was then used to analyze the expression of NK cell activating or inhibitory receptor ligands on GSCs. GSCs (n=6) expressed normal levels of HLA-class I and HLA-E (both ligands for inhibitory NK receptors), at levels similar to those observed on healthy human astrocytes (n=3) **(Fig. 1B)**. In contrast, the ligands for activating NK receptors, such as CD155 (ligand for DNAM1 and TIGIT), MICA/B and ULBP1/2/3 (ligands for NKG2D) and B7-H6 (ligand for NKp30) were upregulated on GSCs but not on healthy human astrocytes **(Fig. 1B)**. To assess the contributions of these activating and inhibitory receptors to the NK cell-dependent cytotoxicity against GSCs, we used receptor-specific blocking antibodies to disrupt specific receptor-ligand interactions. The blockade of NKG2D, DNAM1 and NKp30 but not HLA class I, significantly decreased NK cell-mediated GSC killing (n=4) **(Fig. 1C)**. Cumulatively, these findings suggest that GSCs possess the ligands needed to stimulate NK cell activation leading to GSC elimination. Indeed, the effects we observed were entirely consistent with an extant model of tumor cell attack by NK cells, whereby inhibitory signals transmitted by KIR-HLA class I interactions are overcome when a threshold level of activating signals are reached, inducing recognition of ‘stressed’ cells. ^19, 20, 21, 22^.

**Figure 1.**
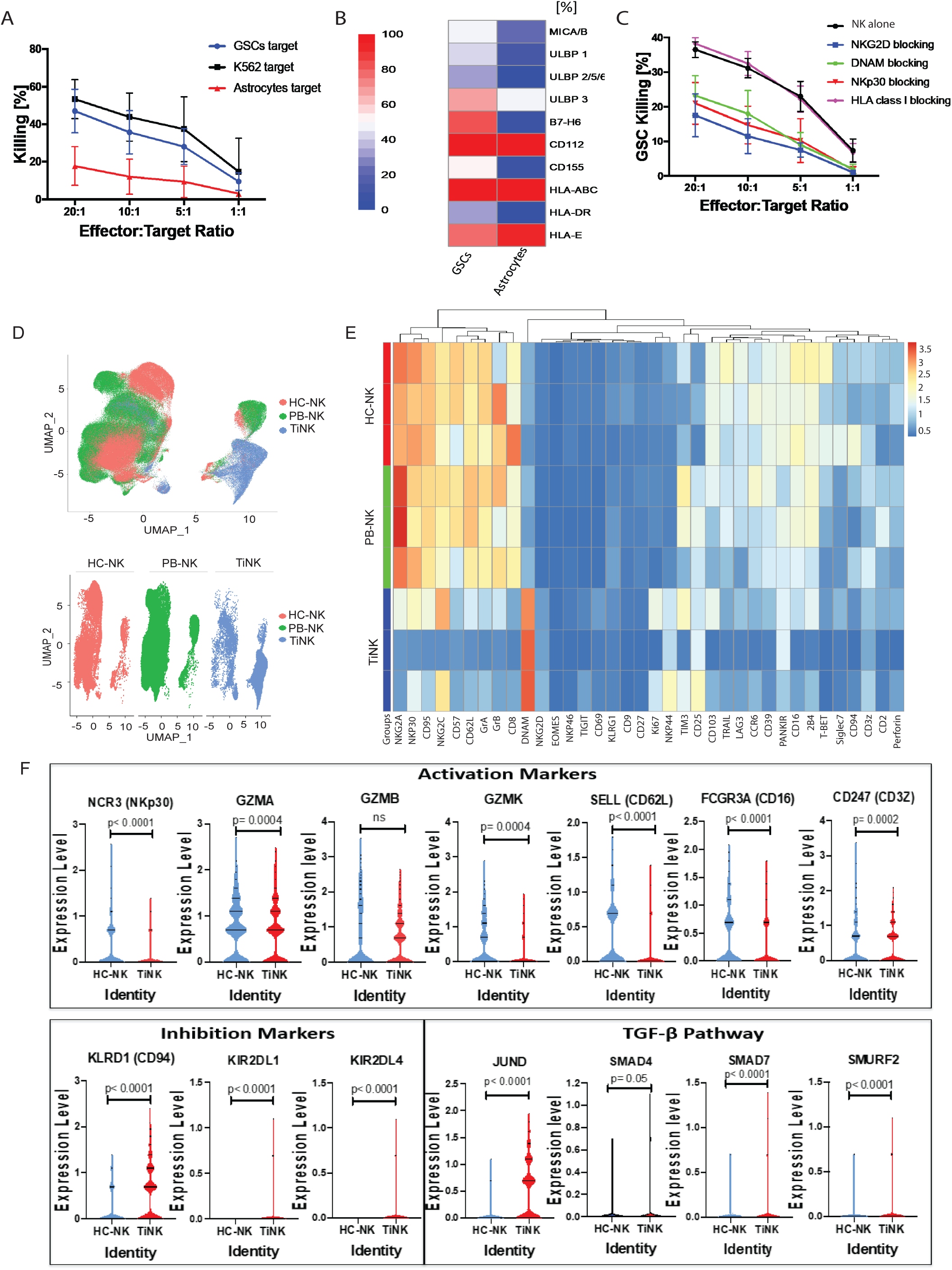
GSCs express NK cell receptor ligands and are susceptible to NK cell cytotoxicity. **A**, Healthy donor-derived NK cells were activated overnight with 5 ng/ml of IL-15 and co-cultured with GBM patient-derived GSCs (blue lines), K562 (black lines) or healthy human astrocytes (red lines) targets for 4 hours at different effector:target ratios. The NK cell cytotoxic activity was measured by ^51^Cr-release assay (n=6). Error bars denote standard deviation. **B**, Summary expression levels of 10 ligands for NK cell receptors on GSCs isolated from GBM patient samples or on healthy human astrocytes. The color scale of the heat map represents the relative expression of NK cell ligand on GSCs or human astrocytes ranging from green (low expression) to red (high expression). The columns present the minimum to maximum and median expression for each receptor (GSC: n=6; Astrocytes: n=3); p=0.03 for ULP2/5/6, p<0.0001 for B7-H6 and p=0.02 for CD155. **C**, Activated NK cells from healthy donors were co-cultured with GSCs for 18 hours in the presence or absence of blocking antibodies against the NK cell receptors NKG2D (blue lines), DNAM (green lines), NKp30 (red lines) or HLA class I (pink lines). The NK cell cytotoxicity against GSCs targets was assessed by ^51^Cr-release assay (n=4). Error bars denote standard error of the mean. **D, E**, viSNE plots (**D**) and a comparative heatmap of mass cytometry data (**E**) showing the expression of NK cell surface markers, transcription factors and cytotoxicity markers in HC-NK (red), GBM patient PB-NK (green) and TiNK (blue). Heatmap column clustering is identified by FlowSOM analysis while each row reflects the expression level for an annotation for an individual patient. Color scale shows the expression level for each marker, with red representing higher expression and blue lower expression (n=3). **F**, Violin plots showing the NK cell mRNA expression levels for individual genes between healthy control PB-NK cells (HC-NK; blue) and TiNK (red) using single-cell RNA sequencing. Markers associated with NK cell activation and cytotoxicity, NK cell inhibition and the TGF-β pathway are presented. P values were derived using unpaired t-test.

### NK cells infiltrate GBM tumors but display an altered phenotype and function

Preclinical findings in glioma-bearing mice indicate that NK cells can cross the blood-brain barrier to infiltrate the brain^23^. However, the limited clinical studies available suggest only minimal NK cell infiltration into GBM tissue^24^. As such, we next investigated whether NK cells are capable of infiltrating into GBMs and their abundance by analyzing *ex vivo* resected glioma tumor specimen collected from 21 of 46 patients with primary or recurrent GBM, and 2 of 5 patients with low-grade gliomas (**Table 1**). The patient characteristics are summarized in **Table 1**. Each gram of GBM contained a median of 166,666 NK cells (range 9,520-600,000; n=21) whereas there were only 500-833 NK cells/g in low-grade gliomas (n=2). These findings indicate that NK cells can traffic into the GBM microenvironment in numbers that appear to be much larger in high-grade gliomas.

**Table 1.** Patient characteristics.

**GBM patients:**
| Patient number | Sex | Age at DOS | Histology | NK cell count/gram tissue | IDH1 status | MGMT status | TP53 status | EGFR status | Previous treatment | Assay utilization |
| --- | --- | --- | --- | --- | --- | --- | --- | --- | --- | --- |
| 1 | M | 57 | pGBM | unknown | unknown | unknown | unknown | unknown | none | phenotype, functional(C+L) |
| 2 | M | 46 | pGBM | 600,000 | neg | unknown | pos | unknown | none | phenotype, functional(C+L) |
| 3 | M | 54 | rGBM | 280,000 | unknown | pos | unknown | unknown | RT+TMZ | phenotype, functional(C+L) |
| 4 | M | 45 | pGBM | 9,520 | neg | unknown | pos | pos | none | phenotype, functional(C) |
| 5 | M | 66 | pGBM | 370,000 | neg | pos | pos | unknown | none | phenotype, functional(C+L) |
| 6 | M | 38 | rGBM | unknown | pos | intermediate | pos | unknown | RT+TMZ | phenotype, functional(C), p-smad |
| 7 | M | 32 | rGBM* | unknown | pos | unknown | pos | unknown | Accutane, XRT+TMZ | Phenotype, p-smad |
| 8 | F | 80 | pGBM | unknown | unknown | unknown | unknown | unknown | none | phenotype, functional(C), p-smad |
| 9 | M | 62 | pGBM | 200,000 | neg | unknown | pos | unknown | none | phenotype, functional(L), p-smad |
| 10 | F | 51 | pGBM | 200,000 | neg | pos | pos | pos | none | phenotype, p-smad |
| 11 | M | 56 | pGBM | unknown | neg | neg | pos | unknown | none | phenotype |
| 12 | F | 30 | rGBM | 300,000 | pos | pos | pos | unknown | XRT+TMZ | Phenotype, p-smad |
| 13 | F | 55 | pGBM | unknown | neg | unknown | pos | pos | none | phenotype |
| 14 | F | 64 | pGBM | 166,666 | neg | unknown | pos | unknown | none | phenotype |
| 15 | M | 31 | pGBM | 133,333 | neg | pos | unknown | unknown | none | functional(L) |
| 16 | F | 43 | pGBM | 100,000 | pos | pos | pos | pos | none | phenotype |
| 17 | M | 60 | pGBM | unknown | neg | neg | unknown | pos | none | phenotype |
| 18 | M | 67 | pGBM | unknown | neg | neg | unknown | pos | none | phenotype, functional(C), reversal |
| 19 | F | 42 | pGBM | unknown | neg | pos | pos | pos | none | phenotype |
| 20 | M | 54 | rGBM | unknown | neg | neg | pos | unknown | RT+TMZ | phenotype |
| 21 | M | 54 | rGBM | unknown | neg | unknown | pos | pos | RT+TMZ | functional(C), reversal |
| 22 | M | 73 | pGBM | 266,666 | neg | pos | pos | pos | none | phenotype, reversal |
| 23 | M | 56 | rGBM | unknown | unknown | unknown | unknown | unknown | XRT+ RT+ Lomustin and avastin | p-smad |
| 24 | F | 50 | pGBM | unknown | neg | unknown | unknown | unknown | none | phenotype |
| 25 | M | 62 | pGBM | 166,666 | unknown | unknown | unknown | unknown | none | reversal |
| 26 | F | 60 | pGBM | unknown | neg | pos | unknown | pos | none | phenotype |
| 27 | F | 42 | pGBM | unknown | unknown | neg | pos | pos | none | reversal |
| 28 | M | 56 | pGBM | 116,666 | neg | neg | pos | pos | none | functional(C) |
| 29 | M | 71 | pGBM | unknown | neg | neg | pos | pos | none | functional(L) |
| 30 | M | 68 | rGBM | unknown | neg | neg | pos | pos | RT+ TMZ | functional(L) |
| 31 | M | 71 | pGBM | unknown | neg | pos | pos | pos | none | phenotype |
| 32 | M | 50 | pGBM | 350,000 | neg | neg | neg | unknown | none | phenotype |
| 33 | F | 61 | pGBM | 250,000 | neg | unknown | pos | unknown | none | p-smad |
| 34 | M | 42 | pGBM | 200,000 | neg | neg | pos | neg | none | p-smad |
| 35 | M | 50 | rGBM | 40,000 | neg | neg | neg | neg | RT+ TMZ | phenotype, p-smad |
| 36 | F | 40 | pGBM | 166,666 | pos | pos | pos | pos | none | reversal |
| 37 | F | 65 | pGBM | 35,000 | neg | pos | pos | pos | none | phenotype |
| 38 | M | 64 | pGBM | 142,857 | neg | neg | pos | pos | none | phenotype |
| 39 | F | 31 | rGBM | 60,000 | neg | neg | pos | neg | RT+TMZ | phenotype |
| 40 | M | 65 | pGBM | unknown | neg | unknown | neg | unknown | none | scRNA seq |
| 41 | M | 53 | pGBM | unknown | neg | unknown | pos | unknown | none | scRNA seq |
| 42 | M | 52 | pGBM | unknown | neg | unknown | pos | unknown | none | scRNA seq |
| 43 | M | 66 | pGBM | unknown | neg | neg | pos | unknown | none | scRNA seq |
| 44 | F | 70 | pGBM | unknown | neg | unknown | pos | unknown | none | scRNA seq |
| 45 | M | 71 | rGBM | unknown | neg | pos | unknown | pos | surgery +TMZ+RT + enrolled in PVSRIPO | scRNA seq |
| 46 | M | 71 | rGBM | unknown | neg | neg | unknown | unknown | surgery + TMZ + RT | scRNA seq |

Low-grade glioma patients:
| Patient number | Sex | Age at DOS | Histology | NK cell count/gram tissue | IDH1 status | MGMT status | TP53 status | Previous treatment | Assay utilization |
| --- | --- | --- | --- | --- | --- | --- | --- | --- | --- |
| 1 | F | 34 | Low grade oligodendroglioma | 500 | pos | positive | pos | none | none |
| 2 | M | 27 | Diffuse astrocytoma | 833 | neg | unknown | neg | none | none |
| 3 | M | 60 | Low grade oligodendroglioma | unknown | pos | unknown | unknown | none | scRNA seq |
| 4 | F | 45 | Diffuse astrocytoma | unknown | pos | pos | pos | none | scRNA seq |
| 5 | M | 39 | Diffuse astrocytoma | unknown | pos | neg | pos | none | scRNA seq |

To gain insights into the phenotype of the GBM tumor-infiltrating NK cells (TiNKs), we used cytometry by time-of-flight (CyToF) and a panel of 37 antibodies against inhibitory and activating receptors, as well as differentiation, homing and activation markers **(Supplementary Table 1)**. We ran uniform manifold approximation and projection (UMAP) a dimensionality reduction method, on a dataset from paired peripheral blood NK cells (PB-NK) and TiNKs from patients with GBM and peripheral blood from healthy controls. Heatmap was used to compare protein expression between the groups. This analysis identified 4 main clusters **(Fig. 1D-E)**. While PB-NK from GBM patient and PB from healthy control (HC-NK) showed great phenotypic similarity, they were markedly different than TiNKs, with the latter characterized by increased expression of CD56^bright^ and significantly lower levels of activating receptors (CD16, NKG2D, NKp30, NKp46, DNAM-1, NKG2C, CD2, CD3z and 2B4), transcription factors (T-bet, eomes), signal transducing adaptor proteins (DAP10, DAP12, SAP) and cytotoxic molecules (granzyme B and perforin) **(Fig. 1E)**. These data were confirmed by multi-parameter flow cytometry in TiNKs and paired PB NK cells from 28 GBM patients and PB samples from 15 HC **(Supplementary Fig. 1A-C)**.

Next, we investigated the NK cell transcriptomic profiles of TiNKs from 10 additional glioma patients and PBMCs from healthy donors using a Drop-Seq-based scRNA-seq technology (10× Genomics <u>STAR Methods</u>) from a soon to be publicly available dataset of CD45+ glioma infiltrating immune cells [Zamler et al. Immune landscape of genetically-engineered murine models of glioma relative to human glioma by single-cell sequencing *Manuscript in Submission.* (2020)]. We analyzed over 1746 NK cells from each GBM patient sample and over 530 cells from each healthy PBMC donor. The NK signature used to define the NK population included the markers KLRD1, NKG7 and NKTR. There was significant downregulation of the genes that encoded NK cell activation markers such as *NCR3* [NKp30], *GZMA* [granzyme A], *GZMK* [granzyme K], *SELL* [CD62L], *FCGR3A* [CD16] and *CD247* [CD3Z] on TiNKs from GBM patients compared with healthy donor PBMCs (HC-NK) **(Fig. 1F)**. Genes that encoded for NK cell inhibitory receptors such as *KLRD1* [CD94], *KIR2DL1* and *KIR2DL4* were upregulated on the TiNKs compared to the HC-NKs **(Fig. 1F)**. Interestingly, genes associated with TGFβ pathway as *JUND, SMAD4, SMAD7* and *SMURF2* were also significantly upregulated on TiNKs compared with HC-NK **(Fig. 1F)**.

We next tested the impact of our phenotypic findings on NK cell function by isolating NK cells from the GBM tumor or PB-NK cells and testing their effector function against K562 targets. TiNKs exerted less cytotoxicity by ^51^Cr release assay, less degranulation (reduced expression of CD107a) and produced significantly lower amounts of IFN-γ and TNF-α than did PB- or HC-NK **(Fig. 2A-B; Supplementary Fig. 2)**. Taken together, these data indicate that NK cells can indeed migrate into GBMs and undergo immune alteration within the tumor microenvironment that results in marked impairment of their cytotoxic function, indicating their susceptibility to immune evasion tactics of the malignant tumor.

**Figure 2.**
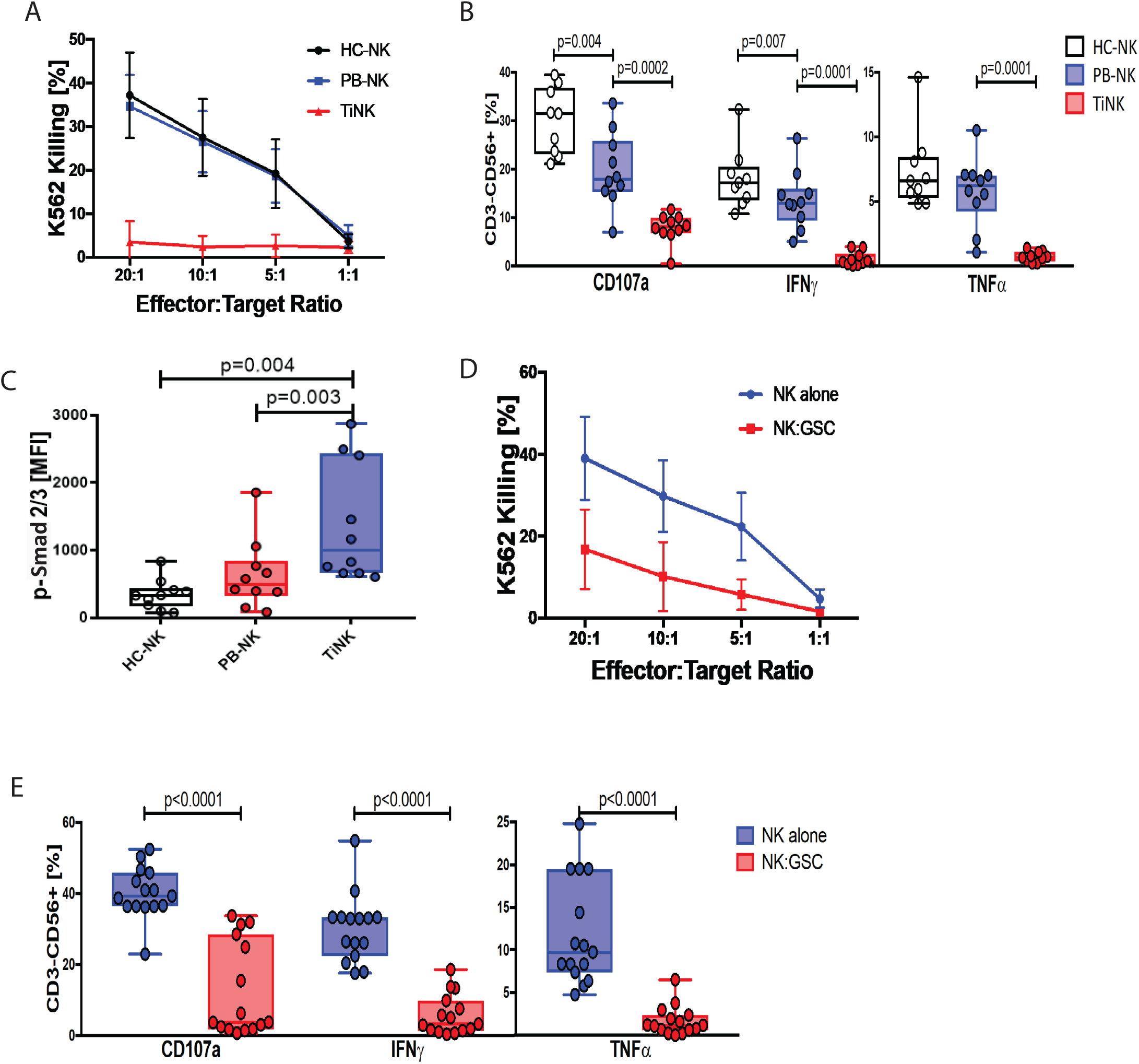
GSCs induce NK cell dysfunction. **A**, Primary human GBM tumor infiltrating NK cells (TiNK) (red lines), paired peripheral blood NK cells (PB-NK) (blue lines) from the same GBM patient or peripheral blood NK cells from healthy control donor (HC-NK) (black lines) were co-cultured for 4 hours with K562 targets at different effector:target ratios and the cytotoxicity was determined by ^51^Cr release assay (n=8). Error bars denote standard deviation. **B**, Box plots summarizing CD107a, IFN-γ, and TNF-α production by TiNKs, PB-NK or HC-NK cells after incubation with K652 targets for 5 hours at a 5:1 effector/target ratio. NK cells were identified as CD3-CD56+ lymphocytes (n=10). Error bars denote standard deviation. Paired t test was performed to determine statistical significance. **C**, Box plots comparison of p-Smad2/3 expression in NK cells from healthy controls (HC-NK, white), PB-NK (red) and TiNKs (blue). Paired t test was performed to determine statistical significance (n=10). **D**, Specific lysis (^51^Cr release assay) of K562 cells by NK cells cultured alone or with GSCs at a 1:1 ratio for 48h. Error bars denote standard deviation (p=0.03, n=10). **E**, Box plots summarizing CD107a, IFN-γ, and TNF-α production by NK cells cultured either alone or with GSCs in a 1:1 ratio for 48 hours in response to K562 targets. Plots are gated on CD3-CD56+ NK cells cultured alone or with GSCs (n=10).

### TGF-β1 mediates NK cell dysfunction in GBM tumors

Despite the intrinsic sensitivity of GSCs to immune attack by NK cells, our findings indicate that this sensitivity is partially lost within the tumor microenvironment, where TiNKs are modulated toward an inhibitory phenotype. Although there are many different mechanisms that could account for this shift in function^12^, the TiNK phenotypic and single cell transcriptomic alterations were most consistent with the effects of TGF-β1, a pleiotropic cytokine that functions as an important inhibitor of the mTOR pathway^25^. This notion was supported by the observation of enhanced basal levels of p-Smad2/3, the canonical TGF-β signaling pathway, in TiNK cells compared to PB- or HC-NK cells **(Fig. 2C; Supplementary Fig. 3)**.

Given the rarity of the GSCs and their exquisite sensitivity to NK cell cytotoxicity, we reasoned that they may have evolved their own mechanisms of immune evasion in addition to the evasive tactics provided by the known immune regulatory cells in the microenvironment^12^. To pursue this hypothesis, we first tested whether GSCs can suppress the function of healthy allogeneic NK cells *in vitro*. While incubation with healthy human astrocytes (control) had no effect on NK cell function (n=3) **(Supplementary Fig. 4A-B)**, co-culture with patient-derived GSCs significantly impaired the ability of allogeneic NK cells to perform natural cytotoxicity and to produce IFN-γ and TNF-α in response to K562 targets (n=10; n=15 respectively) **(Fig. 2D-E; Supplementary Fig. 4C**). Next, we tested whether TGF-β1 plays a role in GSC-induced NK cells dysfunction by co-culturing NK cells from healthy control donors with patient-derived GSCs in the presence or absence of TGF-β neutralizing antibodies and assessing their cytotoxicity against K562 targets. While the antibodies did not affect the normal function of healthy NK cells when cultured alone **(Supplementary Fig. 5A)**, the blockade of TGF-β1 prevented GSCs from disabling NK cell cytotoxicity **(Supplementary Fig. 5B-D)**. Thus, we conclude that TGF-β1 production by GSCs contributes significantly to NK cell dysfunction in the GBM microenvironment.

### Disruption of TGF-β1 signaling prevents but does not reverse GSC-induced NK cell dysfunction

If GSCs induce NK cell dysfunction through activation and release of TGF-β1, it may be possible to avoid this evasive tactic by inhibiting the TGF-β signaling pathway. Thus, we first tested whether galunisertib (LY2157299), a TGF-β receptor I kinase inhibitor that has been used safely in GBM patients^26^, and LY2109761, a dual inhibitor of TGF-β receptors I and II, ^27, 28^ can prevent or reverse GSC-induced NK cell dysfunction. Although neither inhibitor affected NK cell function **(Supplementary Fig. 6A)**, each prevented GSCs from activating the TGF-β1 Smad2/3 signaling pathway in NK cells **(Fig. 3A)** and inducing dysfunction, thus preserving the natural cytotoxicity of NK cells against K562 or GSC targets **(Fig. 3B; Supplementary Fig. 6B-C)**. Interestingly, blockade of the TGF-β receptor kinase by galunisertib or *ex vivo* culture of TiNKs with activating cytokines such as IL-15 failed to inactivate the TGF-β1 Smad2/3 signaling pathway and restore NK cell dysfunction **(Fig. 3C**; **Supplementary Fig. 6D-E**). Similarly, these maneuvers did not reverse the dysfunction of HC-NK cells induced by GSCs **(Supplementary Fig. 6F-G)** indicating that once NK cells are rendered dysfunctional in the suppressive microenvironment of GBM tumors, stimulation with IL-15 or inhibition of TGF-β1 activity is unlikely to restore their function.

**Figure 3.**
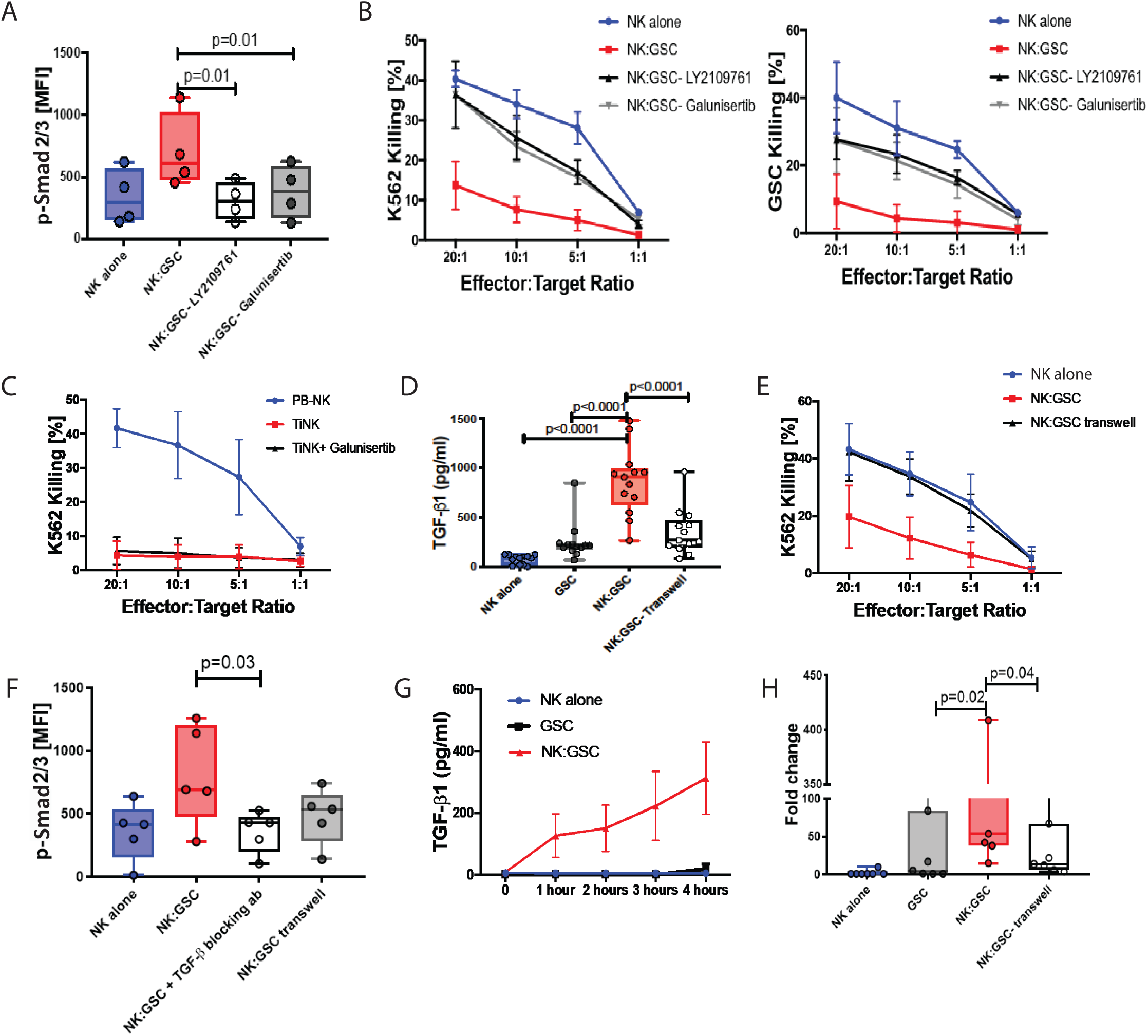
GSC-induced NK cell dysfunction requires cell-to-cell contact. **A**, Box plots summarize the mean fluorescence intensity (MFI) of p-Smad2/3 expression of NK cells cultured alone (NK alone) or co-cultured with GSCs in the presence or absence of the TGF-β receptor small molecule inhibitors LY2109761 (10 *μ*M) and galunisertib (10 *μ*M) for 12 hours. Paired t test was performed to determine statistical significance (n=4). **B**, Specific lysis (^51^Cr release assay) of K562 (left) or GSCs (right) by purified healthy control NK cells after 48 hours of co-culture with or without GSCs in a 1:1 ratio in the presence or absence of LY2109761 (10 *μ*M) (K562: p=0.04; GSC: p=0.02) or galunisertib (10 *μ*M) (K562: p=0.04; GSC: p=0.03) (n=4). Error bars denote standard deviation. **C**, Specific lysis (^51^Cr release assay) of K562 by TiNK after 24 hours of culture either alone or with galunisertib (10 *μ*M) (n=3). **D**, Box plots summarize the soluble TGF-β levels (pg/ml) by Elisa in the supernatant of NK cells and GSCs cultured either alone or together in the presence or absence of a transwell membrane for 48 hours (n=13). P values were derived using unpaired t-test. **E**, Specific lysis (^51^Cr release assay) of K562 by NK cells after 48 hours of co-culture with GSCs in a 1:1 ratio, either in direct contacted or separated by a transwell membrane (p=0.03) (n=7). **F**, Box plots showing the MFI of p-Smad2/3 expression in healthy control NK cells cultured either alone, in direct contact with GSCs in the absence or presence of TGF-β blocking antibodies, or separated from GSCs by a transwell membrane (n=5). P values were derived using unpaired t-test. **G**, Soluble TGF-β levels (pg/ml) in the supernatant of NK cells and GSCs cultured either alone (NK: blue line; GSC: black line) or together (red line) in the first 4 hours of co-culture was measured by ELISA (n=4). **H**, TGF-β mRNA fold change of NK cells and GSCs cultured either alone or together in the presence or absence of a transwell membrane for 48 hours was determined using qPCR (n=7).

### GSCs induce NK cell dysfunction through cell-cell contact dependent TGFβ release

We next asked if TGF-β1 secretion by GSCs is an endogenous process, as observed with macrophages and myeloid-derived suppressor cells (MDSCs)^29, 30^, or requires active cell-cell interaction with NK cells. To address this question, we performed transwell experiments in which healthy donor-derived NK cells and GSCs were either in direct contact with each other or separated by a 0.4 µm pore-sized permeable membrane that allowed the diffusion of soluble molecules, but not cells. Levels of soluble TGF-β1 were measured 48 hours after the cultures were initiated. Direct contact of GSCs with NK cells resulted in significantly higher levels of TGF-β1 compared with those attained when GSCs were separated from NK cells by a transwell (mean 836.9 pg/ml ± 333.1 S.D. vs 349 pg/ml ± 272.2 S.D.) or when GSCs were cultured alone (252 ± 190.4 pg/ml; p<0.0001) **(Fig. 3D)**, indicating that activation and secretion of TGF-β by GSCs is a dynamic process requiring direct cell-cell contact between the NK cells and GSCs. Importantly, healthy human astrocytes cultured either alone or with NK cells did not produce substantial amounts of TGF-β1 **(Supplementary Fig. 7)**. Consistent with these results, we found that GSC-mediated NK cell dysfunction also required direct cell-cell contact. Indeed abrogation of direct cell-cell contact between NK cells and GSCs by a transwell membrane prevented the induction of NK cell dysfunction, and activation of the TGF-β1 Smad2/3 pathway, similar to results with TGF-β1 blocking antibodies **(Fig. 3E-F; Supplementary Fig. 8)**.

TGF-β1 is a tripartite complex and its inactive latent form is complexed with two other polypeptides: latent TGF-β binding protein (LTBP) and latency-associated peptide (LAP). Activation of the mature TGF-β1 requires its dissociation from the engulfing LAP. Because TGF-β1-LAP is expressed on the surface of GSCs at high levels **(Supplementary Fig 9A-B)**, we asked if the increase in soluble TGF-β levels in the supernatant after GSC-NK cell contact was driven by release of the cytokine from the engulfing LAP or by increased transcription of the *TGF-β1* gene, or both. To distinguish between these two alternatives, we investigated if contact with NK cells can induce a rapid release of TGF-β from LAP by measuring the kinetics of TGF-β1 production in the supernatant after GSC-NK cell co-culture. The results indicate a rapid increase in soluble TGF-β1 levels as early as 1 hour after co-culture in conditions where NK cells and GSCs were in direct contact compared with co-cultures in which NK cells and GSCs were cultured alone **(Fig. 3G).** When the fold-changes in TGF-β1 mRNA were determined by quantitative PCR (qPCR) in GSCs alone or in direct contact with NK cells or separated from NK cells by a transwell membrane for 48 hours, the *TGF-β*1 copy numbers were significantly higher in GSCs in direct contact with NK cells (p =0.04) **(Fig. 3H)**. Thus, the marked increase in TGF-β1 seen after NK cell interaction with GSCs appears to involve a dual mechanism of upregulated TGF-β1 transcription and release of the mature cytokine from the LAP peptide by GSCs.

### MMP2 and MMP9 play a critical role in the release of activated TGF-β1 from LAP

Both matrix metalloproteinases (MMPs) 2 and 9 mediate the release of TGF-β1 from LAP^31, 32^. Because both enzymes are expressed by malignant gliomas^33^, we investigated whether they might also be involved in the release of TGF-β1 from LAP and consequently in the induction of NK cell dysfunction by GSCs. First, we confirmed that GSCs are a major source of MMP2 and MMP9 **(Supplementary Fig. 10A-B)**, and then determined their contribution to the release of TGF-β1 and GSC-induced NK cell dysfunction by culturing healthy NK cells with or without GSCs and in the presence or absence of an MMP2/9 inhibitor for 48 hours. MMPs were present at higher levels when GSCs were in direct contact with NK cells, suggesting that TGF-β1 drives their release, as confirmed by experiments using TGF-β blocking antibodies **(Supplementary Fig. 10A-B)**. The addition of an MMP2/9 inhibitor did not affect NK cell function in cultures lacking GSCs **(Supplementary Fig. 10C)** but partially prevented GSC-induced NK dysfunction, as measured by the ability of the NK cells to perform natural cytotoxicity and to produce IFN-γ and TNF-α in response to K562 targets **(Supplementary Fig. 10D-F).** This partial restoration would be consistent with the involvement of additional pathways in the activation of TGF-β. Incubation of NK cells with the MMP2/9 inhibitor also resulted in decreased p-Smad2/3 levels **(Supplementary Fig. 10G)**, implicating MMP2/9 in the release of TGF-β by GSCs.

### αv integrins mediate cell contact dependent TGF-β1 release by GSCs

Since GSC-mediated NK cell dysfunction requires direct cell-cell contact, we next investigated which receptor-ligand interactions could be participating in this crosstalk. Blocking the interaction of major activating and inhibitory NK cell receptors, including CD155/CD112, CD44, KIRs and ILT-2, on healthy donor NK cells and their respective ligands on GSCs failed to prevent GSC-induced NK cell dysfunction **(Supplementary Fig. 11)**. We then changed our focus to the integrins, a family of cell surface transmembrane receptors that play a critical role not only in cell adhesion, migration and angiogenesis, but also in the activation of latent TGF-β1^34^. The αv (CD51) integrin heterodimeric complexes αvβ3, αvβ5 and αvβ8 are highly expressed in glioblastoma, in particular on GSCs^35^. Based on evidence that targeting αv integrins in glioblastoma can significantly decrease TGF-β production,^35^ we tested whether cilengitide, a small molecule inhibitor that possesses a cyclic RDG peptide with high affinity for αv integrins can prevent GSC-induced NK cell dysfunction by decreasing TGF-β1 production. Treatment with cilengitide significantly decreased levels of soluble TGF-β1 **(Fig. 4A)** as well as p-Smad2/3 signaling in NK cells in direct contact with GSCs **(Fig. 4B)** and prevented GSC-induced NK cell dysfunction (n=8; n=12) **(Fig. 4C-E)**. These results were confirmed by genetic silencing of the pan-αv integrin (CD51) in GSCs using CRISPR/Cas9 **(Figure 4F; Supplementary Fig. 12).** Together, our data support a model in which αv integrins regulate the TGF-β1 axis involved in GSC-induced NK cell dysfunction **(Fig. 4G)**.

**Figure 4.**
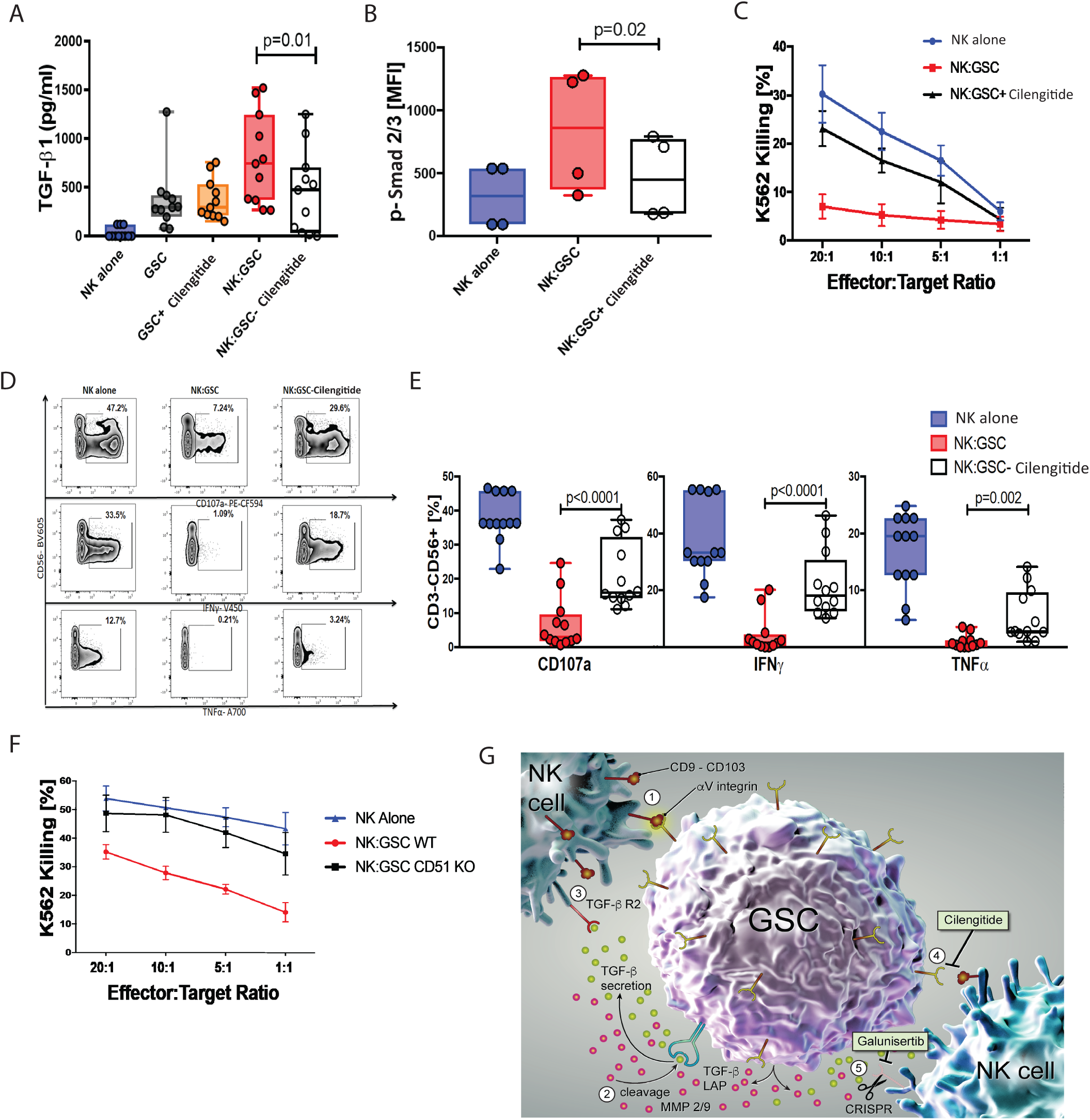
αv integrins mediate TGF-β1 release by GSCs and GSC-induced NK cell dysfunction. **A**, Box plots showing soluble TGF-β levels (pg/ml) in the supernatant of NK cells and GSCs cultured either alone or together in the presence or absence of the αv integrin small molecule inhibitor cilengitide (10 *μ*M) for 48 hours was determined using ELISA (n=11). Error bars denote standard deviation. P values were derived using unpaired t-test. **B**, Box plots showing the MFI of p-Smad2/3 expression on healthy control NK cells cultured either alone or with GSCs in the presence or absence of cilengitide (10 *μ*M). P values were derived using paired t-test. **C**, Specific lysis (^51^Cr release assay) of K562 by NK cells cultured either alone or after co-culture with GSCs for 48 hrs in the presence or absence of cilengitide (10 *μ*M) (p=0.05) (n=8). Error bars denote standard deviation. **D, E**, Representative zebra plots (**D**) and summary box plots (**E**) of CD107, IFN-γ, and TNF-α production by NK cells in response to K562 cultured either alone or after 48 hrs of co-culture with GSCs at a 1:1 ratio with or without cilengitide (n=12). Inset numbers in panel D are the percentages of CD107a-, IFN-γ- or TNF-α-positive NK cells within the indicated regions. **F**, Specific lysis (^51^Cr release assay) of K562 targets by NK cells cultured either alone or with WT GSCs or with *CD51* KO GSCs for 48 hrs at a 1:1 ratio (n=3). Error bars denote standard deviation. **G**, Working hypothesis of GSC-induced NK cell suppression. Cell-cell contact between αv integrins (CD51) on GSCs with surface receptors such as CD9 and CD103 on NK cells mediate the release of TGF-β LAP through shear stress and the release MMP-2/9 from GSCs. Free TGF-β is now able to bind its receptor on NK cells and induce immune suppression. Inhibition of the αv integrins on GSCs or knock out of the TGF-β receptor 2 on NK cells can prevent this chain of events, preserving the cytotoxic activity of NK cells and enabling them to efficiently target GSCs.

We next sought to identify the surface ligands on NK cells that could potentially interact with αv integrins to mediate GSC-NK cell crosstalk. In addition to binding extracellular matrix components, αv integrins bind tetraspanins, such as CD9, through their active RDG binding site^36^. Indeed, CD9 and CD103 are upregulated on GBM TiNKs **(Figure 1E; Supplementary Fig. 1)** and can be induced on healthy NK cells after co-culture with TGF-β1 **(Supplementary Fig. 13A)**. Thus, we used CRISPR Cas9 gene editing to knockout (KO) *CD9* and *CD103* in healthy donor NK cells **(Supplementary Fig. 13B)** and tested the cytotoxicity of wild type (WT, treated with Cas9 only), *CD9* KO, *CD103* KO or *CD9/CD103* double KO NK cells after co-culture with GSCs. As shown in **Supplementary Fig. 13C-E**, silencing of either *CD9* or *CD103* resulted in partial improvement in the cytotoxic function of NK cells co-cultured with GSCs by comparison with WT control. In contrast, CD9/CD103 double KO NK cells co-cultured with GSCs retained their cytotoxicity against K562 targets **(Supplementary Fig. 13C-E)**. This suggests that αv integrins on GSCs bind CD9 and CD103 on NK cells to regulate the TGF-β1 axis involved in GSC-induced NK cell dysfunction.

### Inhibition of the αv integrin TGF-β1 axis enhances NK cell anti-tumor activity *in vivo*

The mechanistic insights gained from the above studies suggest that the αV integrin-TGF-β1 axis regulates an important evasion tactic used by GSCs to suppress NK cell cytotoxic activity and therefore may provide a useful target for immunotherapy of high-grade GBM. To test this prediction, we used a PDX mouse model of patient-derived GSC, in which, ffLuc+ patient-derived GSCs (0.5 x 10^6^) were stereotactically implanted on day 0 through a guide-screw into the right forebrain of NOD/SCID/IL2Rγc null mice (n=4-5 per group). After 7 days, the mice were treated intratumorally with 2.0×10^6^ human NK cells every 7 days for 11 weeks **(Fig. 5A)** with either galunisertib to block the TGF-β signaling or cilengitide to block the integrin pathway. Galunisertib was administered five times a week by oral gavage and cilengitide three times a week by intraperitoneal injection. Animals implanted with tumor that were either untreated or received NK cells alone, galunisertib alone or cilengitide alone served as controls.

**Figure 5.**
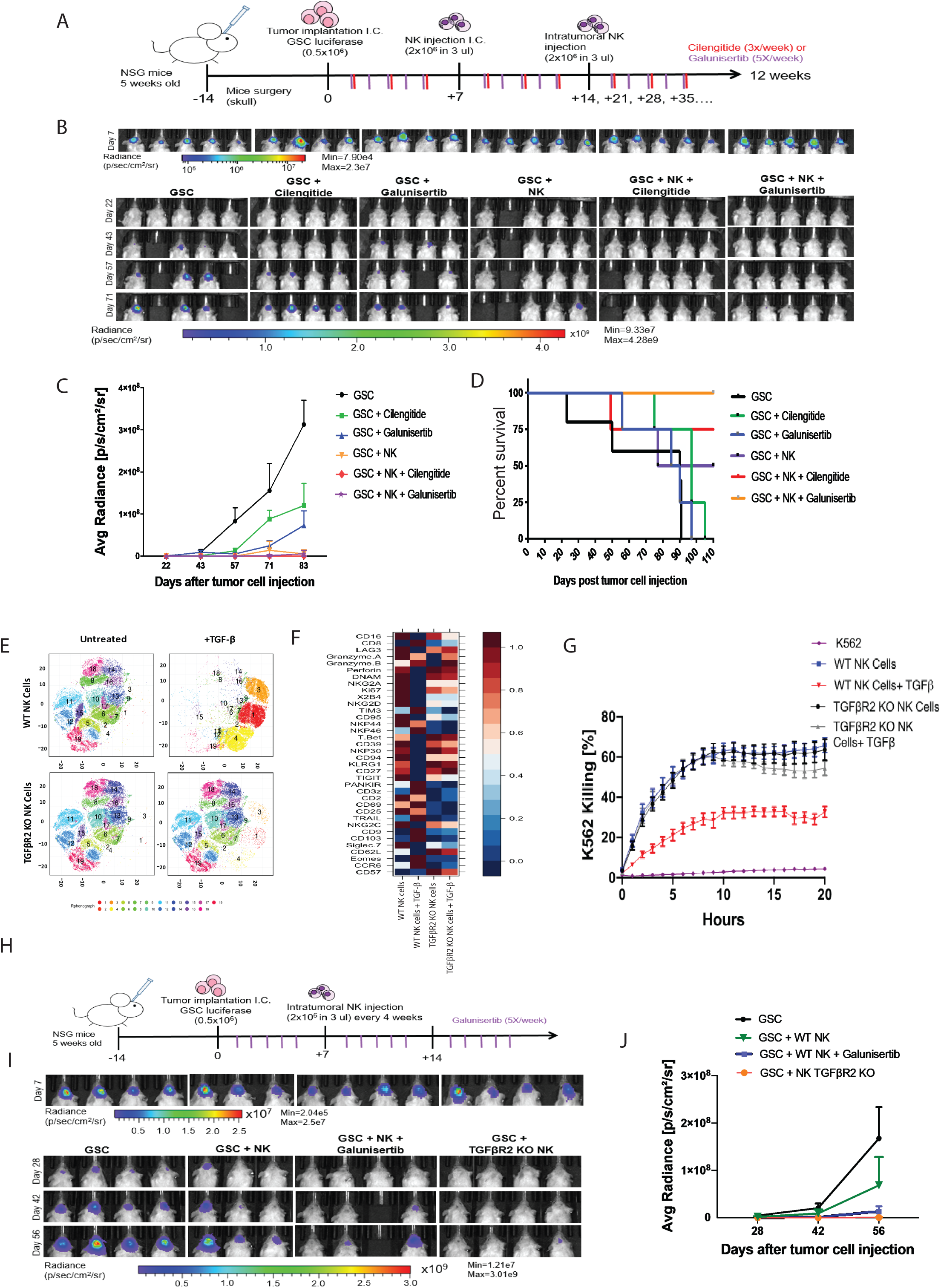
*In vivo* antitumor activity and NK cell function following TGF-β and αv integrin signaling inhibition in NSG GBM mouse model. **A**, Timeline of *in vivo* experiments. GBM tumor implantation was performed at day 0 and *ex vivo*-expanded NK cells were administered intracranially at day 7 and then subsequently every 7 days for 11 weeks. Galunisertib was administered during this time period orally 5 times a week while cilengitide was administered intraperitoneally three times a week for the duration of the experiments. Bioluminescence imaging (BLI) was used to monitor the growth of firefly luciferase-labeled GBM tumor cells in NSG mice. **B**, BLI was obtained from the six group of mice treated with GSC alone (untreated), GSC plus cilengitide, GSC plus galunisertib, GSC plus NK cells, GSC plus NK cells and cilengitide or GSC plus NK cells and galunisertib (4-5 mice per group) as described in panel A. **C**, The plot summarizes the average radiance (BLI) data from our six groups of mice. Mice treated with NK cells together with cilengitide or galunisertib had a significantly lower tumor load by bioluminescence compared with untreated mice or mice treated with cilengitide alone (p<0.0001). Mice treated with NK cells plus galunisertib also had lower tumor load than those treated with galunisertib alone (p=0.04). **D**, Kaplan-Meier plot showing the probability of survival for the groups of mice for each experimental group (5 mice per group). **E, F**, viSNE plots (**E**) and a comparative heatmap (**F**) of mass cytometry data showing the expression of NK cell surface markers, transcription factors and cytotoxicity markers in WT NK cells, *TGFβR2* KO NK cells, WT NK cells + recombinant TGF-β or *TGFβR2* KO NK cells + recombinant TGF-β. Heatmap column clustering, generated by FlowSOM analysis Color scale, shows the expression level for each marker, with red representing higher expression and blue lower expression. **G**, Specific lysis of K562 targets over time by WT-NK (blue), *TGFβR*2 KO (black), WT-NK + recombinant TGF-β (red) or *TGFβR*2 NK cells + recombinant TGF-β (gray) as measured by Incucyte live imaging cell killing assay. **H**, GBM tumor implantation was performed at day 0 and either WT or *TGFβR*2 KO NK cells were administered intracranially at day 7 and then subsequently every 4 weeks. Galunisertib was administered during this time period orally 5 times a week. **I**, BLI was obtained from our three groups of mice: GSCs alone, GSCs plus WT NK cells and galunisertib or GSCs plus *TGFβR*2 KO NK cells (n=4 mice per group). **J**, Plot summarizing the bioluminescence data from our four groups of mice from panel J. Mice treated with WT NK cells and galunisertib or *TGFβR*2 KO NK cells had a significantly lower tumor load by bioluminescence compared with untreated mice (p=0.001; p=0.0002 respectively). Error bars denote standard deviation.

As shown in **Fig. 5B**, tumor bioluminescence rapidly increased in untreated mice and in mice that were either untreated or treated with the monotherapies cilengitide, galunisertib or NK cells. By contrast, adoptive NK cell transfer combined with cilengitide or galunisertib treatment led to significant improvements in tumor control (p<0.0001) **(Fig. 5B-C)** and survival for each comparison (p=0.005) **(Fig. 5D)**. Moreover, no evidence of tissue damage or meningoencephalitis was noted in mice treated with human allogeneic PB-derived NK cells plus cilengitide or galunisertib (**Supplementary Fig. 14**). In animals that received adoptive NK cell infusion combined with either cilengitide or galunisertib, TiNKs harvested after mice were sacrificed showed a higher expression of NKG2D and reduced levels of CD9 and CD103 (**Supplementary Fig. 15**).

Finally, we tested the impact of KO of *TGFβR2* using CRISPR Cas9 gene editing (**Supplementary Fig. 16**) on GSC-induced NK cell dysfunction. *In vitro, TGFβR2* KO NK cells treated with 10 ng/ml of recombinant TGF-β for 48 hours maintained their phenotype compared to wild-type controls as demonstrated by mass cytometry analysis (**Fig. 5E-F**), transcriptomic analysis (**Supplementary Fig. 17A-C**) and cytotoxicity against K562 targets (**Fig. 5G; Supplementary Fig. 17D**). Next, we analyzed the *in vivo* anti-tumor activity of *TGFβR2* KO NK cells by treating mice intracranially at day 7 post tumor implantation with either WT NK cells, WT NK cells plus galunisetib, or *TGFβR2* KO NK cells followed by subsequent NK cell injections every 4 weeks through a guide screw (**Fig. 5H**). Tumor bioluminescence increased rapidly in untreated mice, while adoptive transfer of WT NK cells in combination with 5 x per week galunisertib or *TGFβR2* KO NK cells led to significant tumor control as measured by bioluminescence imaging **(Fig. 5I-J).** In conclusion, our data support a combinatorial approach of NK cell adoptive therapy together with disruption of the αv integrin-TGF-β1 axis to target GBM.

## DISCUSSION

Glioblastoma is among the most deadly and difficult to treat of all human cancers. This difficulty can be in part attributed to the presence of GSCs that differ from their mature progeny in numerous ways, including resistance to standard chemotherapy and radiotherapy, and the ability to initiate tumors and mediate recurrence following treatment. Thus, unless the GSCs within the high-grade GBM tumors are eliminated, the possibility of cure is unlikely. Here, we show that NK cells can readily kill GSCs *in vitro* and can infiltrate these tumors. Yet, they display an altered phenotype with impaired function within the tumor microenvironment, indicating that GSCs have evolved mechanisms to evade NK cell immune surveillance.

Studies to test this hypothesis showed that the underlying mechanism for GSC-induced NK cell dysfunction relies on cell-cell contact between NK cells and GSCs, resulting in the subsequent release and activation of TGF-β, a potent immunosuppressive cytokine that plays a critical role in suppressing the immune response^37^. Our working hypothesis of this protective mechanism is summarized in **Fig. 4G.** Based on our results, we speculate that disruption of the blood-brain barrier caused by the tumor allows the migration of NK cells into the GBM tumor tissue. Once in the tumor, NK cells interact with GSCs. This results in the release and production of TGF-β by GSCs in a cell-cell contact-dependent manner through interactions between αv integrins on GSCs and various ligands on NK cells such as CD9 and CD103. TGF-β is then cleaved from its latent complex form to its biologically active form by proteases such as MMP-2 and MMP-9, released mostly by GSCs. The release of these matrix metalloprotease is further driven by αv integrins and by TGF-β itself, as shown by data presented here and by others ^38, 39, 40, 41, 42, 43, 44^. TGF-β, in turn, suppresses NK cell function by inducing changes in their phenotype, transcription factors, cytotoxic molecules and chemokines. These modifications render NK cells irreversibly incapable of killing GSCs.

An important aspect of this model is the cross-talk between the αv integrins on GSCs and the TGFβ-induced receptors CD9 and CD103 on NK cells as the main mediators of TGF-β production and subsequent NK cell dysfunction. Silencing the pan-αv integrin (CD51) in GSCs by CRISPR/Cas9 gene editing or pharmacologic inhibition with cilengitide prevented GSC-induced NK cell dysfunction, diminished Smad2/3 phosphorylation and decreased TGF-β production in co-cultures of GSCs and NK cells. The αv integrins have been proposed to modulate latent TGF-β activation through two different mechanisms: (i) an MMP-dependent mechanism based on the production of MMP2 and MMP9 by glioma cells and GSCs, but not healthy brain tissue^33^, which proteolytically cleave TGF-β from LAP and (ii), an MMP-independent mechanism, that relies on cell traction forces^38, 41, 43, 44^. This duality may explain why the MMP-2/9 inhibitors used in this study could only partially protect NK cells from GSC-induced dysfunction. Current therapeutic strategies such as radiation therapy may in fact potentiate this vicious cycle of immune evasion. Indeed, radiation therapy has been shown to promote the growth of the therapy resistant GSCs by upregulating TGF-β and integrin expression^45^. Inhibition of the αv integrin-TGF-β axis may thus be crucial not only for the success of immunotherapeutic strategies but also for that of conventional therapies.

Although a number of small molecules that globally inhibit TGF-β are in development for glioblastoma patients, most have been associated with prohibitive toxicity^46^ and lack of efficiency, as shown using trabedersen, a TGF-β2 oligodeoxynucleotide antisense^47^. This could be attributed to the irreversible inhibition of NK cell function through TGF-β released from the GSCs. Since the NK cells have already been adversely affected by the tumor microenvironment, administration of trabedersen would be futile. Given the ubiquitous nature and multiple functions of TGF-β in the central nervous system, we suggest that the use of NK cells to eliminate GSCs within tumor tissues would benefit from concomitant use of αv integrin inhibitors to block TGF-β signaling by GSCs, such as cilengitide that binds αVβ3 and αVβ5 integrins, or gene editing strategies to delete the *TGF-βR2* in NK cells and to protect against TGF-β binding and consequent immunosuppression. Either of these strategies can target local immunosuppressive mechanisms and thus would be expected to reduce excessive toxicity.

Finally, on the strength of these findings, we propose to develop an immunotherapeutic strategy in which third-party NK cells derived from healthy donors are administered in combination with a pan-αv integrin inhibitor or are genetically edited to silence *TGF-βR*2 to protect them from immunosuppression, thus, enabling them to recognize and eliminate rare tumor cells with stem-like properties such as GSCs.

## METHODS

### Patients

Forty-six patients with GBM (n=34 primary GBM; n=12 recurrent GBM) and five patients with low-grade glioma (n=2 low-grade oligodendroglioma; n=3 diffuse astrocytoma) were recruited from The University of Texas MD Anderson Cancer Center (MDACC) for phenotypic (n=28), functional studies (n=14) and single cell RNA sequencing analysis (n=10) (**Table 1**). All subjects gave full informed and written consent under the Institutional Review Board (IRB) protocol number LAB03-0687. All studies were performed in accordance with the Declaration of Helsinki. Buffy coat from normal donors was obtained from gulf Coast Regional Blood Center, Houston, Texas, USA.

### Sample Processing

Peripheral blood mononuclear cells (PBMCs) were purified with Histopaque (Sigma-Aldrich) by density gradient separation. Freshly resected human glioblastoma tissue was minced into small pieces using a scalpel, dissociated using a Pasteur pipette, and suspended in RPMI 1640 medium containing Liberase TM Research Grade Enzyme (Roche) at a final concentration of 30 *μ*g/ml. The prepared mixture was incubated for 1 hour at 37°C with agitation. After brief centrifugation, the pellet was resuspended in 20 ml of 1.03 Percoll (GE Healthcare) underlayed with 10 ml of 1.095 Percoll, and overlayed with 10 ml of 5% FBS in PBS (Hyclone). The tube was centrifuged at 1,200 *g* for 20 minutes at room temperature with no brake. After centrifugation, the cell layer on top of the 1.095 Percoll was collected, filtered through a 70-*μ*m nylon strainer (BD Biosciences), washed and cells were counted using a cellometer (Nexelom Bioscience, Lawrence, MA). NK cells were magnetically purified using NK cells isolation kit (Miltenyi).

### Characterization of GBM tumor infiltrating NK cells (TiNKs), peripheral blood NK cells (PB-NK) and healthy control NK cells (HC-NK)

Flow cytometry: Freshly isolated TiNKs, PB-NK and HC-NK cells were incubated for 20 minutes at room temperature with Live/Dead-Aqua (Invitrogen) and the following surface markers: CD2-PE-Cy7, CD3-APC-Cy7, CD56-BV605, CD16-BV650, NKp30-biotin, DNAM-FITC, 2B4-PE, NKG2D-PE, Siglec-7-PE, Siglec-9-PE, PD-1-BV421, CD103-PE-Cy7, CD62L-PE-Cy7, CCR7-FITC, CXCR1-APC, CX3CR1-PE-cy7, CXCR3-PerCP-Cy5.5 (Biolegend), NKp44-PerCP eflour710 and TIGIT-APC (eBiosciences), streptavidin-BV785, PD-1-V450, CD9-V450 and NKp46-BV711 (BD Biosciences), Human KIR-FITC and NKG2C-APC (R&D), NKG2A-PE-Cy7 and ILT2-APC (Beckman Coulter) and CD57-PerCP (Novus Biological). For detection of intracellular markers, cells were fixed/permeabilized using BD FACS lysing solution and permeabilizing solution 2 according to manufacturer’s instructions (BD Biosciences) followed by intracellular staining with Ki-67-PE and t-bet-BV711 (Biolegend), Eomesodermin-eFluor660 and SAP-PE (eBiosciences), Granzyme-PE-CF594 (BD Biosciences), DAP12-PE (R&D) and DAP10-FITC (Bioss Antibodies) for 30 minutes in room temperature. All data were acquired with BD-Fortessa (BD Biosciences) and analyzed with FlowJo software. The gating strategy for detection of NK cells is presented in **Supplementary Fig. 18.**

### Mass Cytometry

The strategy for antibody conjugation is described elsewhere ^48^. **Supplementary Table 1** shows the list of antibodies used for the characterization of NK cells in the study. Briefly, NK cells were harvested, washed twice with cell staining buffer (0.5% bovine serum albumin/PBS) and incubated with 5 µl of human Fc receptor blocking solution (Trustain FcX, Biolegend, San Diego, CA) for 10 minutes at room temperature. Cells were then stained with a freshly prepared CyTOF antibody mix against cell surface markers as described previously^48, 49^. Samples were acquired at 300 events/second on a Helios instrument (Fluidigm) using the Helios 6.5.358 acquisition software (Fluidigm).

Mass cytometry data were normalized based on EQ™ four element signal shift over time using the Fluidigm normalization software 2. Initial data quality control was performed using Flowjo version 10.2. Calibration beads were gated out and singlets were chosen based on iridium 193 staining and event length. Dead cells were excluded by the Pt195 channel and further gating was performed to select CD45+ cells and then the NK cell population of interest (CD3-CD56+). A total of 320,000 cells were proportionally sampled from all samples to perform automated clustering. The mass cytometry data were merged together using Principal Component Analysis (PCA), “RunPCA” function, from R package Seurat (v3). Dimensional reduction was performed using “RunUMAP” function from R package Seurat (v3) with the top 20 principal components. The UMAP plots were generated using the R package ggplot2 (v3.2.1). Data were analyzed using automated dimension reduction including (viSNE) in combination with FlowSOM for clustering ^50^ for the deep phenotyping of immune cells as published before ^51^. We further delineated relevant cell clusters, using our in-house pipeline for cell clustering. To generate the heatmap, CD45+CD56+CD3-gated FCS files were exported from FlowJo to R using function “read.FCS” from the R package flowCore (v3.10). The markers expression was transformed using acrsinh with a cofactor of 5. The mean values of 36 markers were plotted as heat map using the function “pheatmap” from R package pheatmap (v1.0.12). Markers with similar expression were hierarchically clustered.

### Incucyte live imaging

After co-culture with GSCs, NT NK cells and *TGFβRII* KO NK cells were purified and labeled with Vybrant DyeCycle Ruby Stain (ThermoFisher) and co-cultured at a 1:1 ratio with K562 targets labeled with CellTracker Deep Red Dye (ThermoFisher). Apoptosis was detected using the CellEvent Caspase-3/7 Green Detection Reagent (ThermoFisher). Frames were captured over a period of 24 hrs at 1 hour intervals from 4 separate 1.75 x 1.29 mm^2^ regions per well with a 10× objective using IncuCyte S3 live-cell analysis system (Sartorius). Values from all four regions of each well were pooled and averaged across all three replicates. Results were expressed graphically as percent cytotoxicity by calculating the ratio of red and green overlapping signals (count per image) divided by the red signal (counts per image).

### Single cell RNA sequencing

Gliomas were mechanically dissociated with scissors while suspended in Accutase solution (Innovative Cell Technologies, Inc.) at room temperature and then serially drawn through 25-, 10- and 5-mL pipettes before being drawn through an 181/2-gauge syringe. After 10 minutes of dissociation, cells were spun down at 420 x g for 5 minutes at 4°C and then resuspended in 10 mL of a 0.9N sucrose solution and spun down again at 800 x g for 8 minutes at 4°C with the brake off. Once sufficient samples were accumulated to be run in the 10x pipeline (10x Genomics; 6230 Stoneridge Mall Road, Pleasanton, CA 94588), cells were then thawed and resuspended in 1 mL of PBS containing 1% BSA, for manual counting. Cells were then stained with the CD45 antibody (BD Biosciences, San Jose, CA, cat #: 555482) at 1:5 for 20 minutes on ice. Samples had Sytox blue added just before sorting so that only live CD45+ cells would be collected. Cells were then sorted in a solution of 50% FBS and 0.5% BSA in PBS, spun down, and resuspended at a concentration of 700-1200 cells/*μ*L for microfluidics on the 10x platform (10x Genomics). The 10x protocol, which is publicly available, was followed to generate the cDNA libraries that were sequenced. (https://assets.ctfassets.net/an68im79xiti/2NaoOhmA0jot0ggwcyEKaC/fc58451fd97d9cbe012c0abbb097cc38/CG000204_ChromiumNextGEMSingleCell3_v3.1_Rev_C.pdf). The libraries were sequenced on an Illumina next-seq 500, and up to 4 indexed samples were multiplexed into one output flow cell using the Illumina high-output sequencing kit (V2.5) in paired-end sequencing (R1, 26nt; R2, 98nt, and i7 index 8nt) as instructed in the 10x Genomics 3’ Single-cell RNA sequencing kit. The data were then analyzed using the cellranger pipeline (10x Genomics) to generate gene count matrices. The mkfastq argument (10x Genomics) was used to separate individual samples with simple csv sample sheets to indicate the well that was used on the i7 index plate to label each sample. The count argument (10x Genomics) was then used with the expected number of cells for each patient. The numbers varied between 2,000 and 8,000 depending on the number of viable cells isolated. Sequencing reads were aligned with GRCh38. The aggr argument (10x Genomics) was then used to aggregate samples from each patient for further analysis. Once gene-count matrices were generated, they were read into an adapted version of the Seurat pipeline^19,20^ for filtering, normalization, and plotting. Genes that were expressed in less than three cells were ignored, and cells that expressed less than 200 genes or more than 2500 genes were excluded, to remove potentially poor- and high-PCR artifact cells, respectively. Finally, to generate a percentage of mitochondrial DNA expression and to exclude any cells with more than 25% mitochondrial DNA (as these may be doublets or low-quality dying cells), cells were normalized using regression to remove the percent mitochondrial DNA variable via the scTransform^21^ command which corrects for batch effects as well. Datasets were then processed for principal component analysis (PCA) with the RunPCA command, and elbow plots were printed with the ElbowPlot command in order to determine the optimal number of PCs for clustering; 15 PCs were chosen for this analysis. Next, the cell clusters were identified and visualized using SNN and UMAP, respectively, before generating a list of differentially-expressed genes for each cluster. A list of differentially-expressed genes was generated to label our clusters at low resolution (0.1). These clusters’ labels were based on at least three differentially-expressed genes, and violin plots were generated to show the relative specificity to the cluster. Differentially-expressed genes were identified using cutoffs for min.pct = 0.25 and logfc.threshold = 0.25. Plots were generated with either the DimPlot, FeaturePlot or VlnPlot commands. Next we identified the clusters containing NK cell populations in both the PBMC and GBM dataset: NK markers included KLRD1, NKG7, and NKTR. Analyses performed on the combination of PBMC and GBM NK cells, were joined using the FindIntegrationAnchors command to determine genes that can be used to integrate the two datasets—after the determination of the Anchors we used the IntegrateData command to combine our two datasets. Data were then normalized using the scTransform command. Datasets were then processed for PCA with the RunPCA command, and elbow plots were printed with the ElbowPlot command in order to determine the optimal number of PCs for clustering, 15 PCs were chosen for this analysis. Next, the cell clusters were identified and visualized using SNN and UMAP, respectively, before generating a list of differentially-expressed genes for each sample. Plots were generated with either the DimPlot, FeaturePlot or VlnPlot commands.

### GSC culture

GSCs were obtained from primary human GBM samples as previously described^52, 53^. Patients gave full informed and written consent under the IRB protocol number LAB03-0687. The GSCs were cultured in stem cell-permissive medium (neurosphere medium): Dulbecco’s Modified Eagle Medium containing 20 ng/ml of epidermal growth factor and basic fibroblast growth factor (all from Sigma-Aldrich), B27 (1:50; Invitrogen, Carlsbad, CA), 100 units/ ml of penicillin and 100 mg/ml streptomycin (Thermo Fisher Scientific, Waltham, MA) and passaged every 5–7 days ^54^. All generated GSC cell lines used in this paper were generated at MD Anderson Cancer Center and referred to as MDA-GSC.

### NK cell expansion

NK cells were purified from PBMCs from healthy donors using an NK cell isolation kit (Miltenyi Biotec, Inc., San Diego, CA, USA). NK cells were stimulated with irradiated (100 Gy) K562-based feeder cells engineered to express 4-1BB ligand and CD137 ligand (referred to as Universal APC) at a 2:1 feeder cell : NK ratio and recombinant human IL-2 (Proleukin, 200 U/ml; Chiron, Emeryville, CA, USA) in complete CellGenix GMP SCGM Stem Cell Growth Medium (CellGenix GmbH, Freiburg, Germany) on day 0. After 7 days of expansion, NK cells were used for *in vivo* mice experiments and for *in vitro* studies.

### Characterization of GSCs and human astrocytes

Human fetal astrocytes cell lines were purchased from Lonza (CC-2565) and Thermo Fisher Scientific (N7805100) and the human astroglia cell line (CRL-8621) was purchased from the American Type Culture Collection (ATCC). The cells were separated into single cell suspension using accutase (Thermo Fisher Scientific) for GSCs and trypsin for the attached astrocytes. The cells were then stained for MICA/B-PE, CD155-PE-Cy7, CD112-PE, HLA-E-PE and HLA-ABC-APC (Biolegend), ULBP1-APC, ULBP2/5/6-APC and ULBP3-PE (R&D), HLA-DR (BD Biosciences) and B7-H6-FITC (Bioss antibodies) for 20 minutes before washing and acquiring by flow cytometry.

### NK cell cytotoxicity assay

NK cells were co-cultured for 5 hours with K562 or GSCs target cells at an optimized effector:target ratio of 5:1 together with CD107a PE-CF594 (BD Biosciences), monensin (BD GolgiStopTM) and BFA (Brefeldin A, Sigma Aldrich). NK cells were incubated without targets as the negative control and stimulated with PMA (50 ng/mL) and ionomycin (2 mg/mL, Sigma Aldrich) as positive control. Cells were collected, washed and stained with surface antibodies (mentioned above), fixed/permeabilized (BD Biosciences) and stained with IFN-γ v450 and TNF-α Alexa700 (BD Biosciences) antibodies.

### Chromium release assay

NK cell cytotoxicity was assessed using chromium (^51^Cr) release assay. Briefly, K562 or GSCs target cells were labeled with ^51^Cr (PerkinElmer Life Sciences, Boston, MA) at 50 *μ*Ci/5 × 10^5^ cells for 2 hours. ^51^Cr-labeled K562/GSC targets (5 × 10^5^) were incubated for 4 h with serially diluted magnetically isolated NK cells in triplicate. Supernatants were then harvested and analysed for ^51^Cr content.

### Suppression assay

For studies of NK cell suppression by GSCs and human astrocytes, magnetically selected healthy NK cells were cultured in Serum-free Stem Cell Growth Medium (SCGM; CellGro /CellGenix) supplemented with 5% glutamine, 5 *μ* M HEPES (both from GIBCO/ Invitrogen), and 10% FCS (Biosera) in 96-well flat-bottomed plates (Nunc) at 100,000/100*μ*l. NK cells were co-cultured either alone (positive control) or with GSCs or astrocytes at a 1:1 ratio for 48 hours at 37 °C before performing functional assays to assess NK cells cytotoxicity.

### Functional assays for blocking NK cytotoxicity

Magnetically purified NK cells were cultured alone or with blocking antibodies against NKG2D (clone 1D11), DNAM (clone 11A8) and NKp30 (clone P30-15) (Biolegend) overnight (5 *μ*g/ml). ^51^Cr release assay was then performed as described above. For HLA-KIR blocking, GSCs were cultured alone or with an HLA-ABC blocking antibody (clone W6/32, Biolegend) before performing ^51^Cr release assay.

### NK cell functional assays

GSCs and purified NK cells were co-cultured for 48 hours in the presence of anti–TGFβ 123 (5 *μ*g/ml) (R&D), HLA-ABC blocking antibody (clone W6/32, Biolegend), CD44 blocking antibody (clone IM7, Biolegend), ILT-2 (CD85J) blocking antibody (clone HP-F1, ThermoFisher), CD155 blocking antibody (clone D171, GenTex), CD112 blocking antibody (clone TX31, Biolegend), 10 *μ*M LY2109761, 10 *μ*M galunisertib (LY2157299), 10 *μ*M Cilengitide (Cayman Chemical) or 1 *μ*M MMP-2/MMP-9 inhibitor I (Millipore). Cytotoxicity assays were then performed as described above.

### Transwell assays

NK cells (1x 10^5^) were either added directly to GSCs at a ratio of 1:1 or placed in transwell chambers (Millicell, 0.4 *μ*m; Millipore) for 48 hours at 37°C. After 48 hours, cultured cells were harvested to measure NK cell cytotoxicity by both ^51^Cr release assay and cytokine secretion assay.

### NK cell recovery assays

NK cells were cultured either with GSCs in a 1:1 ratio or alone for 48 hours. After 48 hours of co-incubation, NK cells were then either purified again by bead selection and resuspended in SCGM media or remained in culture with GSCs for an additional 48 hours and then used for ^51^Cr release assay. In a second assay, after reselection, NK cells were cultured for another 5 days in SCGM in the presence of 5 ng/ml IL-15 with or without 10 *μ*M galunisertib before use for ^51^Cr release assay.

### TGF-β ELISA and MMP2/9 luminex

NK cells and GSCs were either co-cultured or cultured alone for 48 hours in serum free SCGM growth medium. After 48 hours, supernatants were collected and secretion of TGF *β* and MMP2/3/9 was assessed in the supernatant by TGF*β*1 ELISA kit (R&D systems) or MMP2/3/9 luminex kit (eBiosciences) as per the manufacturer’s protocol.

### Reverse transcriptase–polymerase chain reaction (RT-PCR), and quantitative real-time PCR (qPCR)

RNA was isolated using RNeasy isolation kit (Qiagen). A 1 µg sample of total RNA was reverse transcribed to complementary DNA using the iScript cDNA Synthesis Kit (Bio-Rad) according to the manufacturer’s instructions. Then, an equivalent volume (1 µL) of complementary DNA (cDNA) was used as a template for quantitative real-time PCR (qPCR) and the reaction mixture was prepared using iTaq™ Universal SYBR® Green Supermix (Biorad) according to the manufacturer’s instructions. Gene expression was measured in a StepOnePlus™ (Applied Biosystem) instrument according to the manufacturer’s instruction with the following gene-specific primers: *TGFB1* (forward, 5’-AACCCACAACGAAATCTATG-3’; reverse, 5’-CTTTTAACTTGAGCCTCAGC-3’); and *18S* (forward, 5’- AACCCGTTGAACCCCATT -3’; reverse, 5’-CCATCCAATCGGTAGTAGCG-3’). The gene expression data were quantified using the relative quantification (ΔΔCt) method, and 18S expression was used as the internal control.

### CRISPR gene editing of primary NK cells and GSCs

crRNAs to target CD9, CD103 and CD51 were designed using the Integrated DNA Technologies (IDT) predesigned data set. Guides with the highest on-target and off-target scores were selected. The crRNA sequences are reported in **Supplementary Table 2**. crRNAs were ordered from IDT (www.idtdna.com/CRISPR-Cas9) in their proprietary Alt-R format. Alt-R crRNAs and Alt-R tracrRNA were re-suspended in nuclease-free duplex buffer (IDTE) at a concentration of 200 µM. Equal amount from each of the two RNA components was mixed together and diluted in nuclease-free duplex buffer at a concentration of 44 µM. The mix was boiled at 95°C for 5 minutes and cooled down at room temperature for 10 minutes. For each well undergoing electroporation, Alt-R Cas9 enzyme (IDT cat # 1081058, 1081059) was diluted to 36 µM by combining with resuspension buffer T at a 3:2 ratio. The guide RNA and Cas9 enzyme were combined using a 1:1 ratio from each mixture. The mixture was incubated at room temperature for 10–20 minutes. Either a 12 well plate or a 24 well plate was prepared during the incubation period. This required adding appropriate volume of media and Universal APCs (1:2 ratio of effector to target cells) supplemented with 200 IU/ml of IL-2 (for NK cells only) into each well. Target cells were collected and washed twice with PBS. The supernatant was removed as much as possible without disturbing the pellet and the cells were resuspended in Resuspension Buffer T for electroporation. The final concentration for each electroporation was 1.8 µM gRNA, 1.5 µM Cas9 nuclease and 1.8 µM Cas9 electroporation enhancer. The cells were electroporated using Neon Transfection System, at 1600V, 10ms pulse width and 3 pulses with 10ul electroporation tips (Thermo Fisher Scientific (cat # MPK5000)). After electroporation the cells were transferred into the prepared plate and placed in the 37C incubator. The knockout efficiency was evaluated using flow cytometry 7 days after electroporation. Anti-CD51-PE antibody (Biolegend) was used to verify KO efficacy in GSCs.

To knockout *TGFβR*2, two sgRNA guides (**Supplementary Table 2**) spanning close regions of exon 5 were designed and ordered from IDT; 1 *μ*g cas9 (PNA Bio) and 500 ng of each sgRNA were incubated on ice for 20 minutes. After 20 minutes, NK cells 250,000 were added and re-suspended in T-buffer to a total volume of 14ul (Neon Electroporation Kit, Invitrogen) and electroporated before transfer to culture plate with APCs as described above.

### Phospho-Smad2/3 assay

NK cells were stained with Live/dead-aqua and CD56 ECD (Beckman Coulter) for 20 min in the dark at RT, washed with PBS fixed for 10 min in the dark. After one wash, the cells were permeabilized (Beckman Coulter kit) and stained with p-(S465/S467)-Smad2/p- (S423/S425)/Smad3-Alexa 647 mAb Phosflow antibody (BD Biosciences) for 30 minutes at room temperature. Cells incubated with 10ng/ml recombinant TGF-β for 45 minutes in 37°C were used as positive control.

### MMP2 and MMP9 intracellular staining and western blotting

NK cells and GSCs were either cultured alone, in a transwell chamber or together in the presence or absence of TGF-β blocking antibodies (R&D) for 48 hours. BFA was added for the last 12 hours of culture. Cells were then fixed/permeabilized (BD Biosciences) and stained with anti-MMP2-PE (R&D) and MMP9-PE (Cell Signalling) for 30 minutes before acquisition of data by flow cytometry. The surface markers CD133, CD3 and CD56 were used to distinguish NK cells and GSCs for data analysis.

### Xenogeneic mouse model of GBM

To assess the anti-tumor effect of NK cells against GSCs *in vivo*, we used a NOD/SCID IL-2Rγnull (NSG) human xenograft model (Jackson Laboratories, Bar Harbor, ME). Intracranial implantation of GSCs into male mice was performed as previously described^55^. A total of 60 mice were used. 0.5 × 10^5^ GSCs were implanted intracranially into the right frontal lobe of 5 week old NSG mice using a guide-screw system implanted within the skull. To increase uniformity of xenograft uptake and growth, cells were injected into 10 animals simultaneously using a multiport Microinfusion Syringe Pump (Harvard Apparatus, Holliston, MA). Animals were anesthetized with xylazine/ketamine during the procedure. For *in vivo* bioluminescent imaging, GSCs were engineered to express luciferase by lentivirus transduction. Kinetics of tumor growth was monitored using weekly bioluminescence imaging (BLI; Xenogen-IVIS 200 Imaging system; Caliper, Waltham, MA). Signal quantitation in photons/second (p/s) was performed by determining the photon flux rate within standardized regions of interest (ROI) using Living Image software (Caliper). 2×10^6^ in 3 *μ*l expanded donor peripheral blood NK cells^56^ were injected intracranially via the guide-screw at day 7 post tumor implantation, and then every 7 days for 11 weeks. Mice were treated with either cilengitide or galunisertib (both from MCE Med Chem Express, Monmouth Junction, NJ) in the presence or absence of intracranial NK cell injection. Cilengitide was administered intraperitoneally 3 times a week starting at day 1 (250 *μ*g/100 *μ*l PBS) while galunisertib was administered orally (75 mg/kg) by gavage 5 days a week starting at day 1 (see **Figure 5A**). In a second experiment, mice were injected intracranially via the guide screw 7 days post tumor inoculation with either wild type (WT) NK cells, WT NK cells plus galunisertib or *TGFβR2* KO NK cells followed by subsequent NK cells injections every 4 weeks as describe above. Mice that presented neurological symptoms (i.e. hydrocephalus, seizures, inactivity, and/or ataxia) or moribund were euthanized. Brain tissue was then extracted and processed for NK cells extraction. All animal experiments were performed in accordance with recommendations in the Guide for the Care and Use of Laboratory Animals of the National Institute of Health, and approved by the Institutional Animal Care and Use Committee (IACUC) protocol number 00001263-RN01 at MD Anderson Cancer Center.

### Mice brain tissue processing and analysis

Brain tissue from the animals was collected and NK cells were isolated using a percoll (GE Healthcare) gradient fallowing protocol described by Pino et al ^57^. Briefly, brain tissue was dissociated using a 70 µm cell strainer (Life Science, Durham, NC). Cell suspension was re suspended in a 30% isotonic percoll solution and layer on top of a 70% isotonic percoll solution. Cells were centrifuged at 500G for 30 minutes and 18°C with no brake. 2-3 ml of the 70%-30% interface was collected in a clean tube and washed with PBS 1X. After this procedure, cells were ready for immunostaining with mouse CD45, human CD45, CD56, CD3, CD103, CD9, CD69, PD-1 and NKG2D all from Biolegend.

### Histopathology

Brain tissue specimens from untreated control mice, mice treated with either NK cells alone, cilengitide alone, galunisertib alone or with combination therapy of NK + cilengitide or NK + galunisertib were collected. The specimens were bisected longitudinally and half of each brain was fixed in 10% neutral buffered formalin and were then embedded in paraffin. Formalin-fixed, paraffin embedded tissues were sectioned at 4 μm, and stained routinely with hematoxylin and eosin. Brains were examined for the presence or absence of glioblastoma tumor cells. Sections lacking tumor were also evaluated for evidence of meningoencephalitis using a Leica DM 2500 light microscope by a board-certified veterinary pathologist. One section was examined from each sample. Representative images were captured from comparable areas of cerebral hemispheres with a Leica DFC495 camera using 1.25x, 5x, and 20x objectives.

### Statistical analyses

Statistical significance was assessed with the Prism 6.0 software (GraphPad Software, Inc.), using unpaired or paired two-tailed t-tests as appropriate. For survival comparison a Log-rank test was used. Graphs represent mean and standard deviation (SD). For comparison of bioluminescence among the treatment groups ANOVA was used. A P ≤ 0.05 was considered to be statistical significance.

## Acknowledgments

We thank our summer students Nadia Agha (University of Houston), Cindy Saliba (University of Iowa) and Lihi Shalev (Hebrew University of Jerusalem) for their assistance with some of the experiments performed in the paper. This work was supported in part by the Dr. Marnie Rose Foundation and the MD Anderson Cancer Center Glioblastoma Moonshot, part of the institution’s Moon Shots Program, by a grant (CA016672) to the MD Anderson Cancer Center from the NIH and by Specialized Program of Research Excellence (SPORE) in Brain Cancer grant (P50CA127001). The MD Anderson Flow Cytometry and Cellular Imaging Core Facility (FCCICF), NCI Cancer Center Support Grant (P30CA16672) assisted with the CyTOF studies in this project.

## AUTHOR CONTRIBUTIONS

HS, MHS and RB performed experiments, interpreted and analyzed data. KG, JG, AA, NU, SL, JJL, SA, EG, JY, NWF, LL, MK, MB, AKNC, ELE, DZ, ALG and LNK assisted with experiments and commented on the manuscript. KC, FW, QM, JD, YS and VM performed statistical analysis and commented on the manuscript. KG and CK provided clinical data. MD, JW, MM, LL, MM, EJS, PB, EL, DY, RC, MB, SM, GO, NI, ME, MK, JH, GD and FL provided advice on experiments and commented on the manuscript. KR and AH designed and directed the study. HS, RB, KR, DM, MHS and LMF wrote the manuscript.

## CONFLICT OF INTERESTS

The authors have no relevant conflict of interests to declare.

## DATA AVAILABILITY

The data that support the findings of this manuscript are available from the corresponding author upon reasonable request.

