## Supplementary Materials for "Inhibition of the αv integrin-TGF-β axis improves natural killer cell function against glioblastoma stem cells"

Supplementary Figures

Supplementary Figure 1

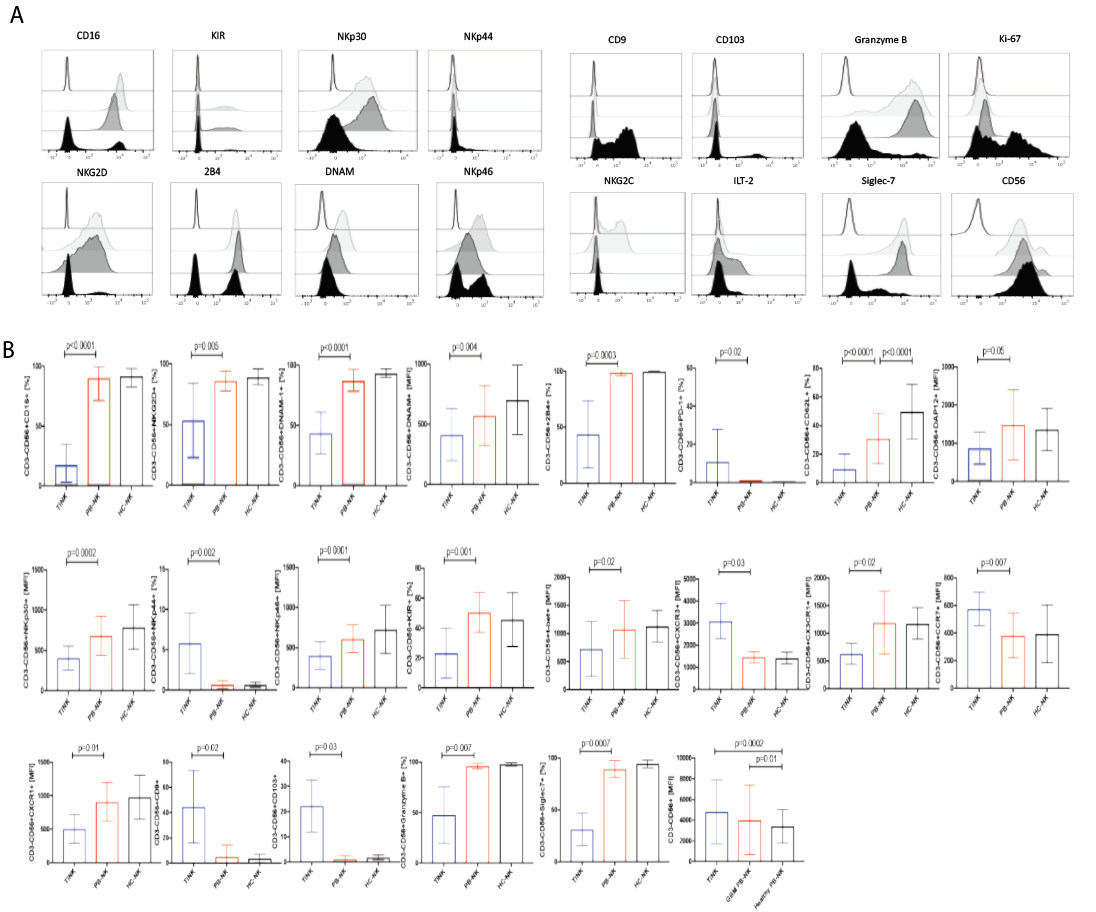

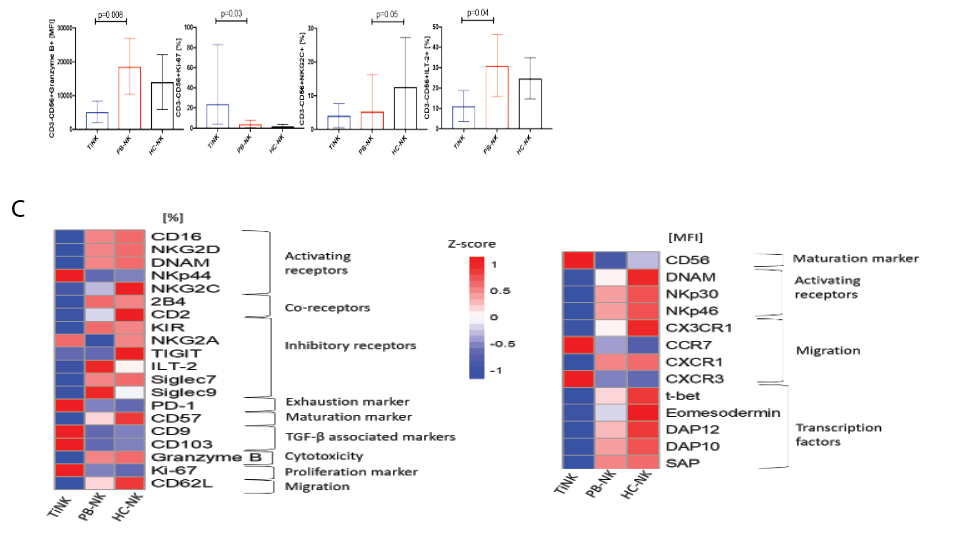

**Supplementary Figure 1. GBM tumor infiltrating NK cells phenotype by flow cytometry. A-C,** Representative histograms and graph summary for the mean fluorescence intensity (MFI) or frequencies of NK cells expressing individual markers in GBM tumor infiltrating NK cells (TiNKs) vs. autologous peripheral blood (PB-NK) from the same GBM patient vs. peripheral blood from healthy controls (HC-NK). Error bars denote mean and standard deviation. P values were derived using paired (PB-NK vs. TiNK) and unpaired (HC-NK vs. TiNK) t-tests (n=28). **C,** The color scale of the heatmap represents the relative expression for each marker ranging from blue (low expression) to red (high expression).

Supplementary Figure 2

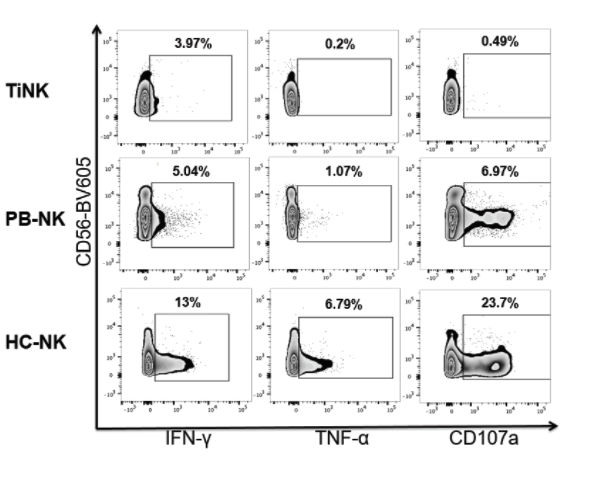

**Supplementary Figure 2. GBM TiNK cells are dysfunctional.** Representative zebra plot for CD107a, IFN-γ, and TNF-α production by TiNKs, PB-NK or HC-NK cells after incubation with K562 targets for 5 hours at a 5:1 effector : target ratio. Inset numbers are the percentages of CD107a-, IFN-γ- or TNF-α-positive NK cells within the gated populations.

Supplementary Figure 3

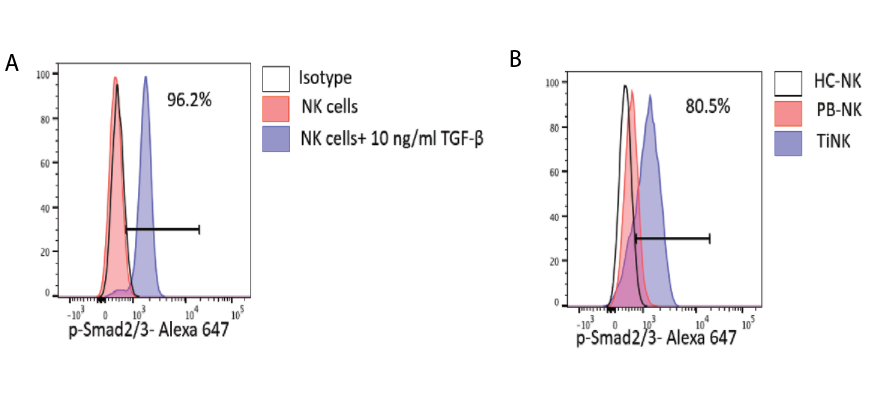

**Supplementary Figure 3. TGF-β induces phosphorylation of Smad2/3 proteins in human NK cells by flow cytometry. A,** Representative histograms show the levels of p-Smad2/3 at baseline (red histogram) and after 30 minutes stimulation with 10 ng/ml of recombinant TGF-β in healthy control NK cells. **B**, Representative histograms show the baseline levels of p-Smad2/3 in healthy control HC-NK cells (white histogram), GBM PB-NK cells (red histogram) and GBM TiNK cells (blue histogram).

Supplementary Figure 4

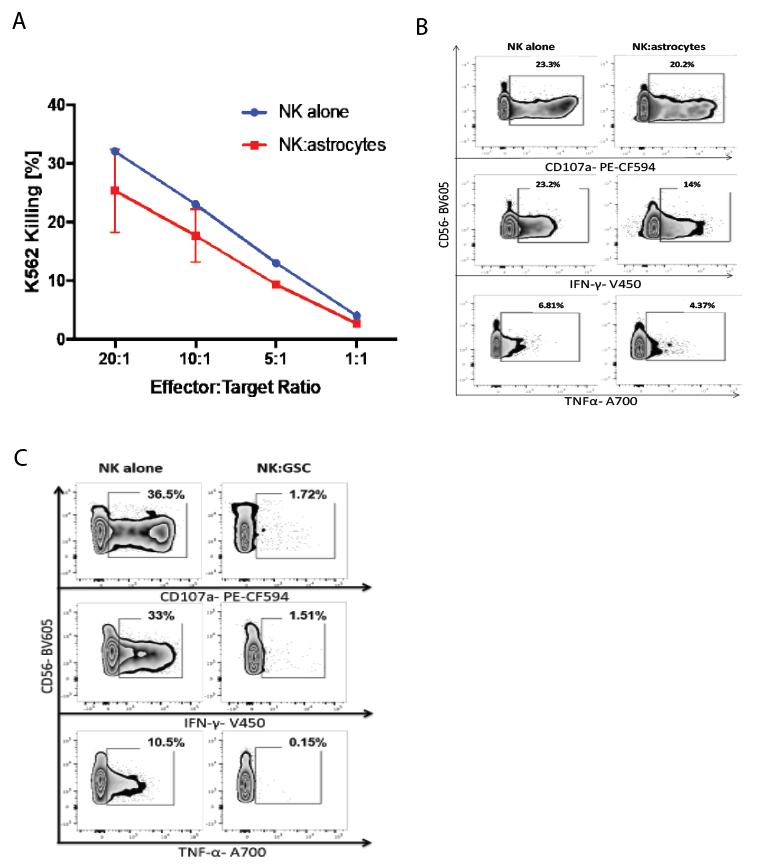

**Supplementary Figure 4. GSCs but not healthy astrocytes induce NK cell dysfunction in vitro. A**, Specific lysis (^51^Cr release assay) of K562 targets by NK cells cultured either alone (blue lines) or with healthy human astrocytes (red lines) in a 1:1 ratio for 48 hours (n=3). **B**, Heathy donor NK cells were co-cultured with astrocytes for 48 hours at a 1:1 ratio. Representative zebra plots show their **CD107a**, IFN-γ, and TNF-α response to K562 targets. Effector:target ratio is 5:1. NK cells were gated on CD3-CD56+ lymphocytes (n=3). Inset numbers are the percentages of CD107a-, IFN-γ- or TNF-α-positive NK cells within the gated populations. **C**, Heathy donor NK cells were co-cultured with GSCs for 48 hours at a 1:1 ratio. Representative zebra plots show their CD107a, IFN-γ, and TNF-α production in response to K562 targets. NK cells were defined as CD3-CD56+ lymphocytes (n=3). Inset numbers are the percentages of CD107a-, IFN-γ- or TNF-α-positive NK cells within the gated populations.

Supplementary Figure 5

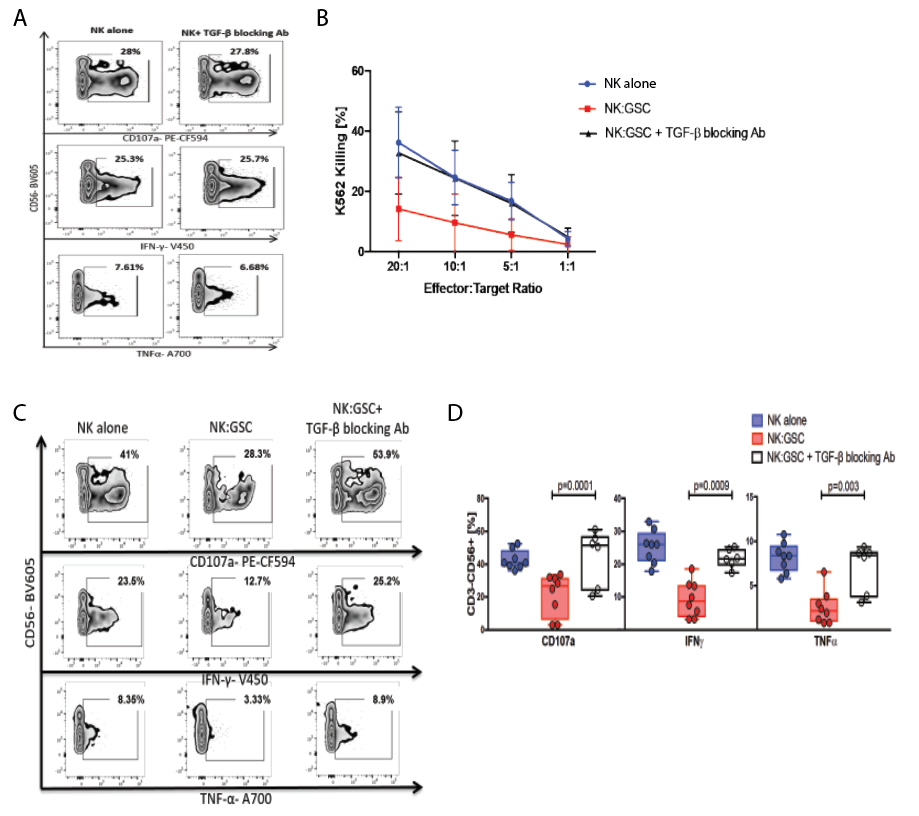

**Supplementary Figure 5. Blockade of TGF-β prevents GSC-induced NK cell dysfunction. A**, Representative zebra plots for CD107a, IFN-γ, and TNF-α production by NK cells in response to K562 targets after incubation with or without TGF-β blocking antibody (5 μg/ml). Effector:target ratio is 5:1. NK cells were gated on CD3-CD56+ lymphocytes. Inset numbers are the percentages of CD107a-, IFN-γ- or TNF-α-positive NK cells within the gated population. **B**, Specific lysis (^51^Cr release assay) of K562 targets by NK cells cultured either alone (blue lines) or co-cultured with GSCs for 48 hours at different effector:target ratios in the presence (black lines) or absence (red lines) of TGF-β blocking antibody (5 μg/ml) (p=0.04) (n=5). **C-D**, Heathy donor NK cells were co-cultured with GSCs at a 1:1 ratio for 48 hours, in the presence or absence of TGF-β blocking antibody (5 μg/ml), Representative zebra plots and summary box plots show their CD107a, IFN-γ, and TNF-α response to K562 targets. Effector:target ratio is 5:1. NK cells were defined as CD3-CD56+ lymphocytes. Inset numbers are the percentages of CD107a-, IFN-γ- or TNF-α-positive NK cells within the gated population (n=5).

Supplementary Figure 6

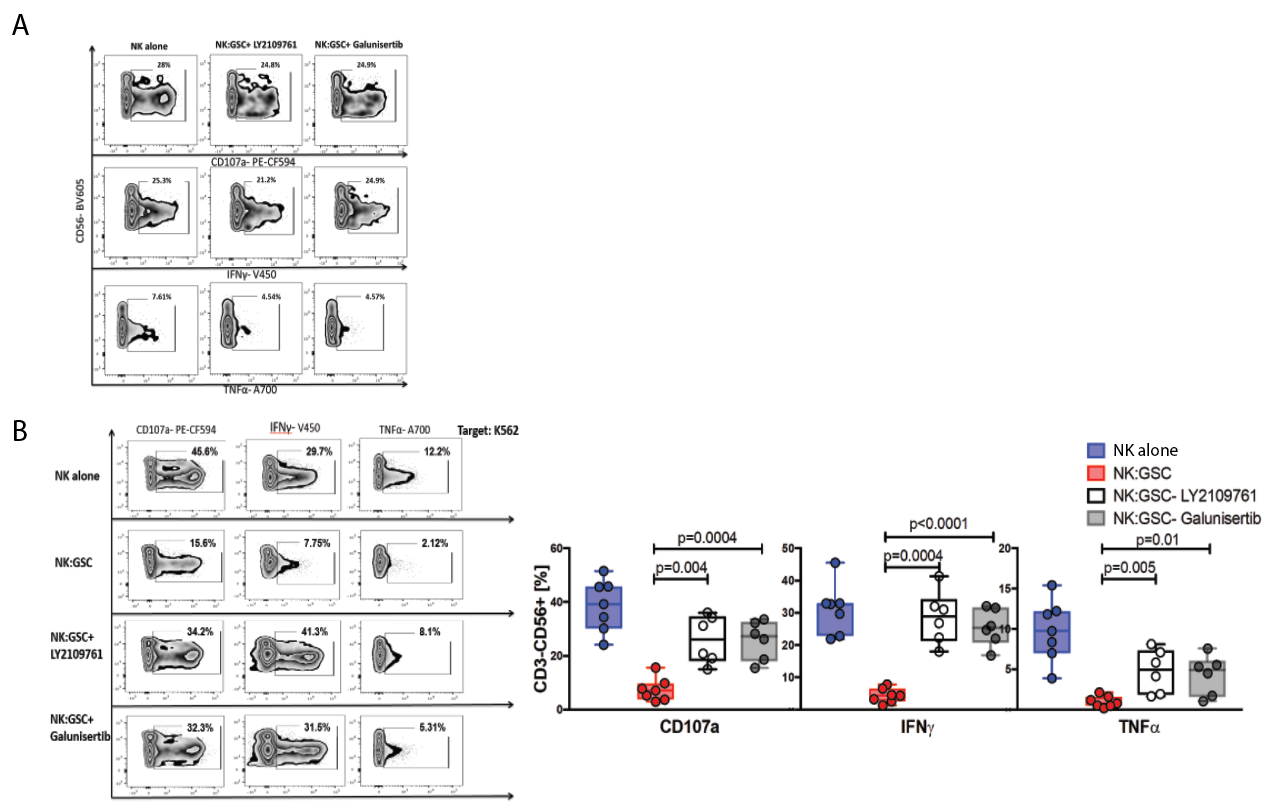

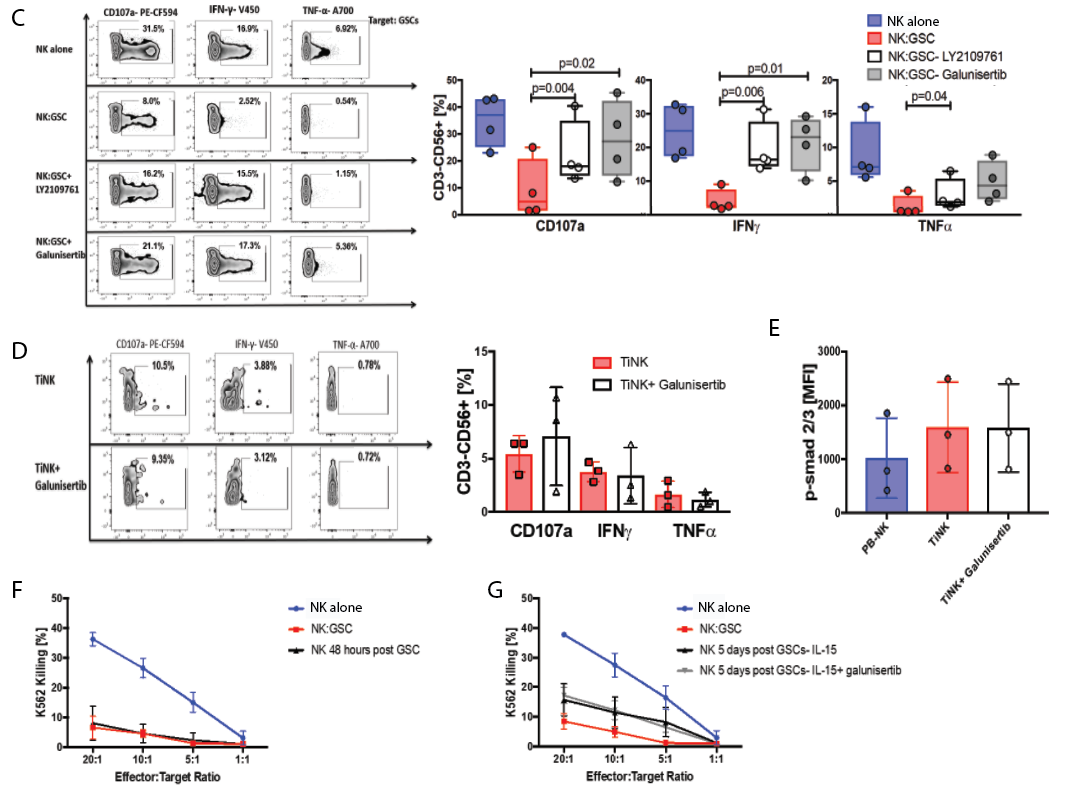

**Supplementary Figure 6. The TGF-β receptor kinase inhibitors Galunisertinib and LY2109761 prevent but do not reverse GSC-induced NK cell dysfunction in vitro**. **A**, NK cells were incubated with or without galunisertinib (10 μM) or LY2109761 (10 μM) for 48 hours. Representative zebra plots show their CD107a, IFN-γ, and TNF-α response to K562 targets. NK cells were gated on CD3-CD56+ lymphocytes. Inset numbers are the percentages of CD107a-, IFN-γ- or TNF-α-positive NK cells within the gated NK cell population. **B**,**C,** NK cells were cultured either alone or with GSCs in a 1:1 ratio with or without LY2109761 or galunisertib for 48 hrs. Representative zebra plots and summary box plots show their CD107, IFN-γ, and TNF-α expression response to K562 (**B**) or GSC (**C**) targets. Effector:target ratio is 5:1.. NK cells were defined as CD3-CD56+ lymphocytes. Inset numbers are the percentages of CD107a-, IFN-γ- or TNF-α-positive NK cells (n=7, n=4 respectively). **D**, TiNK cells were cultured for 24 hours in media with 5 ng/ml IL-15 with or without galunisertib. Representative zebra plots and bar graph summary show their CD107, IFN-γ, and TNF-α response to K562 targets. Effector:target ratio is 5:1. NK cells were gated on CD3-CD56+ lymphocytes. Inset numbers are the percentages of CD107a-, IFN-γ- or TNF-α-positive NK cells within the indicated regions. (n=3). **E**, TiNK cells and paired PB-NK cells from GBM patients were cultured in the presence or absence of galunisertib for 12 hours. The bar graphs summarize the mean fluorescence intensity (MFI) for their p-Smad2/3 expression. Paired t test was performed to determine statistical significance (n=3). **F**, Specific lysis (^51^Cr release assay) of K562 targets by NK cells. After 48 hours of culture alone (blue lines) or with GSCs, healthy donor NK cells were either left with GSCs (red lines), or purified and re-suspended in media for another 48 hours (black lines). NK cells were then collected and used for ^51^Cr release assay (n=4). **G**, Specific lysis (^51^Cr release assay) of K562 targets by NK cells. After 48 hours of culture alone (blue lines) or with GSCs, healthy donor NK cells were either left with GSCs (blue lines) or purified and cultured in SCGM media plus 5 ng/ml IL-15 with (black lines) or without galunisertib (10 μM) (gray lines) for 7 days. At the end of the culture period, their cytotoxicity was tested against k562 targets by ^51^Cr release assay (n=4).

Supplementary Figure 7

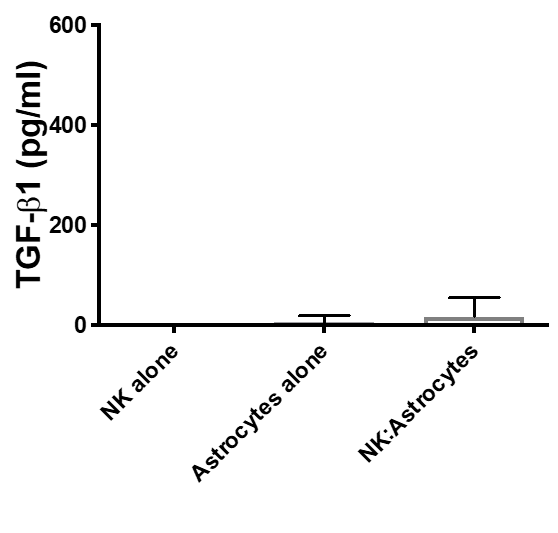

**Supplementary Figure 7. NK cells or astrocytes cultured either alone or together do not produce soluble TGF-β**. Soluble TGF-β1 levels (pg/ml) measured by ELISA in the supernatant of NK cells and astrocytes cultured either alone or in direct contact at a 1:1 ratio for 48 hours (n=3)**.**

Supplementary Figure 8

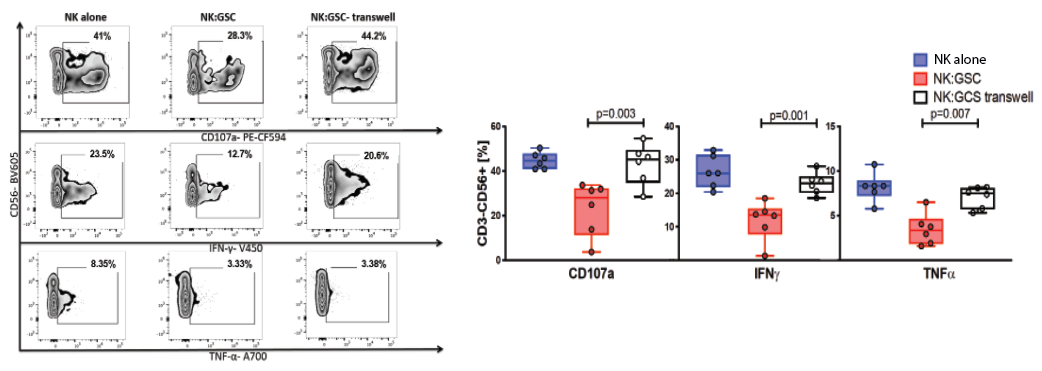

**Supplementary Figure 8. GSC-induced NK cell dysfunction is mediated through cell-cell contact.** Healthy donor NK cells were cultured for 48 hours either alone, or with GSCs (1:1 ratio) in direct contact or with separation by a transwell membrane (n=6). Representative zebra and box plots summarize their CD107a, IFN-γ, and TNF-α response to K562. Effector:target ratio is 5:1. NK cells were gated on CD3-CD56+ lymphocytes. Inset numbers are the percentages of CD107a-, IFN-γ- or TNF-α-positive NK cells within the indicated regions.

Supplementary Figure 9

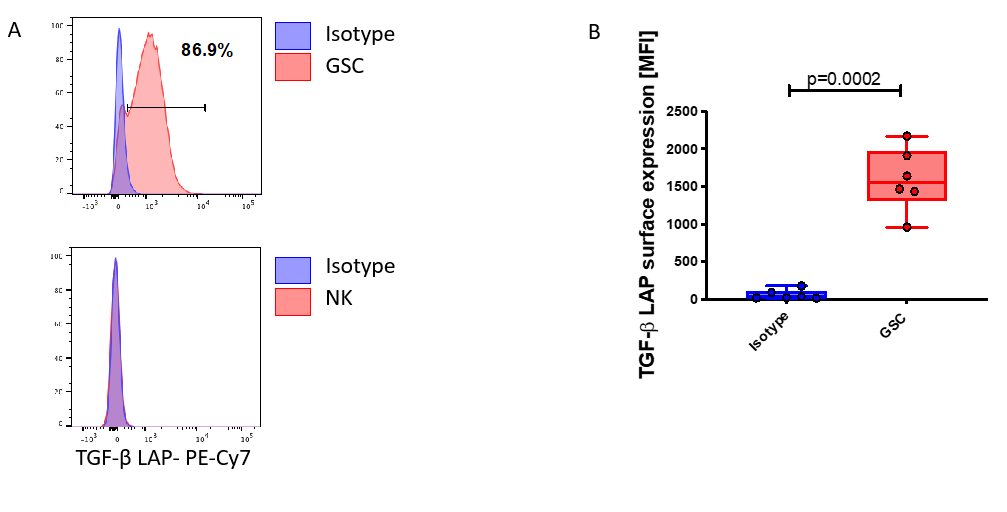

**Supplementary Figure 9. TGF-β latency-associated peptide (LAP) is expressed on the surface of GSCs but not on NK cells. A**, Representative histograms show TGF-β LAP expression on the surface of GSCs and NK cells (blue histogram). Isotype control is shown in red. Inset numbers are the percentages of TGF-β LAP-positive GSCs (top) vs. NK cells (bottom) within the gated population. **B**, Box plots summarize the TGF-β LAP surface expression on GSCs as measured by MFI (n=6). Error bars denote standard deviation. P values were derived using paired t-test.

Supplementary Figure 10

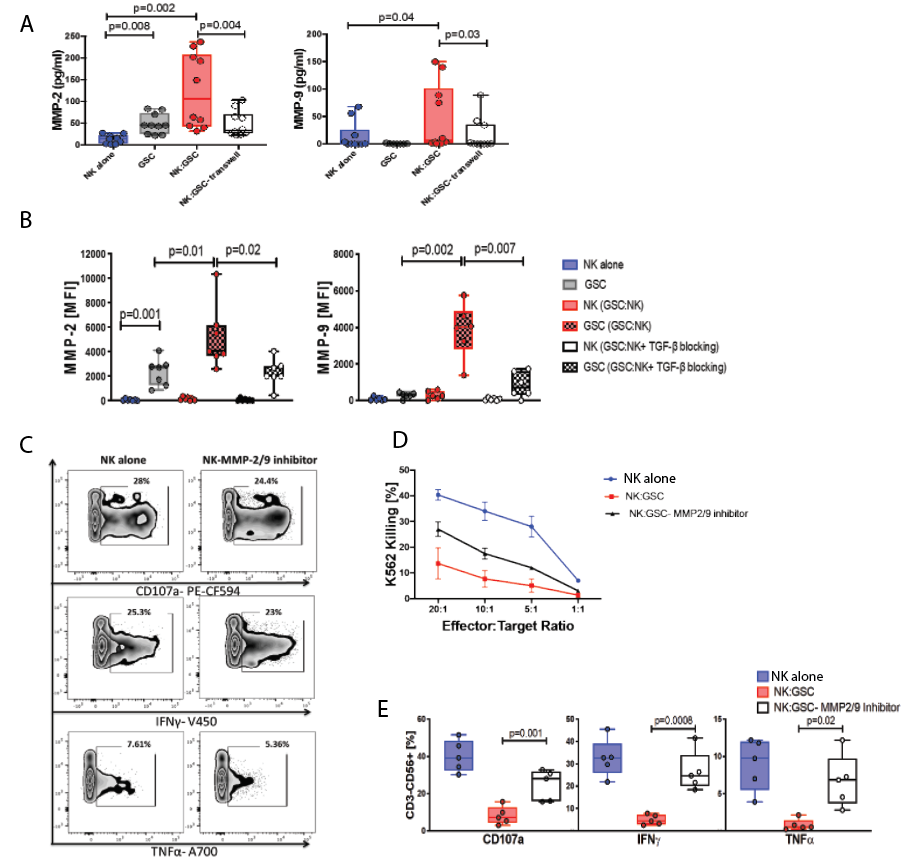

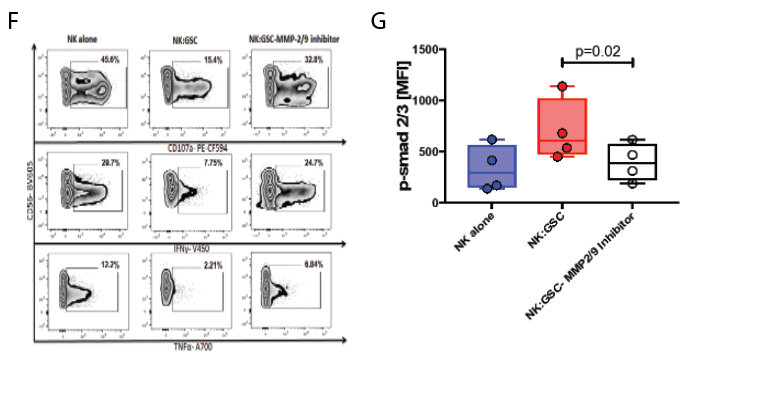

**Supplementary Figure 10. MMP2 and MMP9 partially regulate TGF-β release by GSCs. A,** Box plots summarizing the levels of MMP2 and MMP9 (pg/ml) in the supernatant of NK cells and GSCs cultured either alone or together for 48 hours, either in direct cell contact or separated by a transwell membrane were measured using Luminex assay (n=10). P values were derived using paired t-test. **B**, Box plots showing the MFI for MMP2 and MMP9 expression on NK cells or on GSCs cultured either alone or together in the presence or absence of 5 μg/ml of TGF-β blocking antibody (n=7). P values were derived using paired t-test. **C,** Healthy donor NK cells were cultured with or without an MMP2/9 inhibitor (1 μM) for 48 hours and their CD107a, IFN-γ, and TNF-α response to K562 targets was measured. NK cells were gated on CD3-CD56+ lymphocytes. Inset numbers are the percentages of CD107a-, IFN-γ- or TNF-α-positive NK cells within the indicated regions. **D,** Healthy donor NK cells were cultured either alone (blue lines), or with GSCs at a 1:1 ratio with (black likes) or without (red lines) the MMP2/9 inhibitor (1 μM) for 48 hrs. Their cytotoxicity (^51^Cr release assay) was measured against K562 targets (p=0.04) (n=3). **E, F,** Healthy donor NK cells were cultured either alone or with GSCs at a 1:1 ratio with or without the MMP2/9 inhibitor for 48 hrs. Representative zebra plots and box plots summarize their CD107, IFN-γ, and TNF-α response to K562 targets (n=5). Effector:target ratio is 5:1. Inset numbers are the percentages of CD107a-, IFN-γ- or TNF-α-positive NK cells within the indicated regions. P values were derived using paired t-test. **G,** Box plot show the expression of p-Smad2/3, as measured as MFI in NK cells in the presence or absence of GSCs, with or without the MMP2/9 inhibitor (n=4). P values were derived using paired t-test.

Supplementary Figure 11

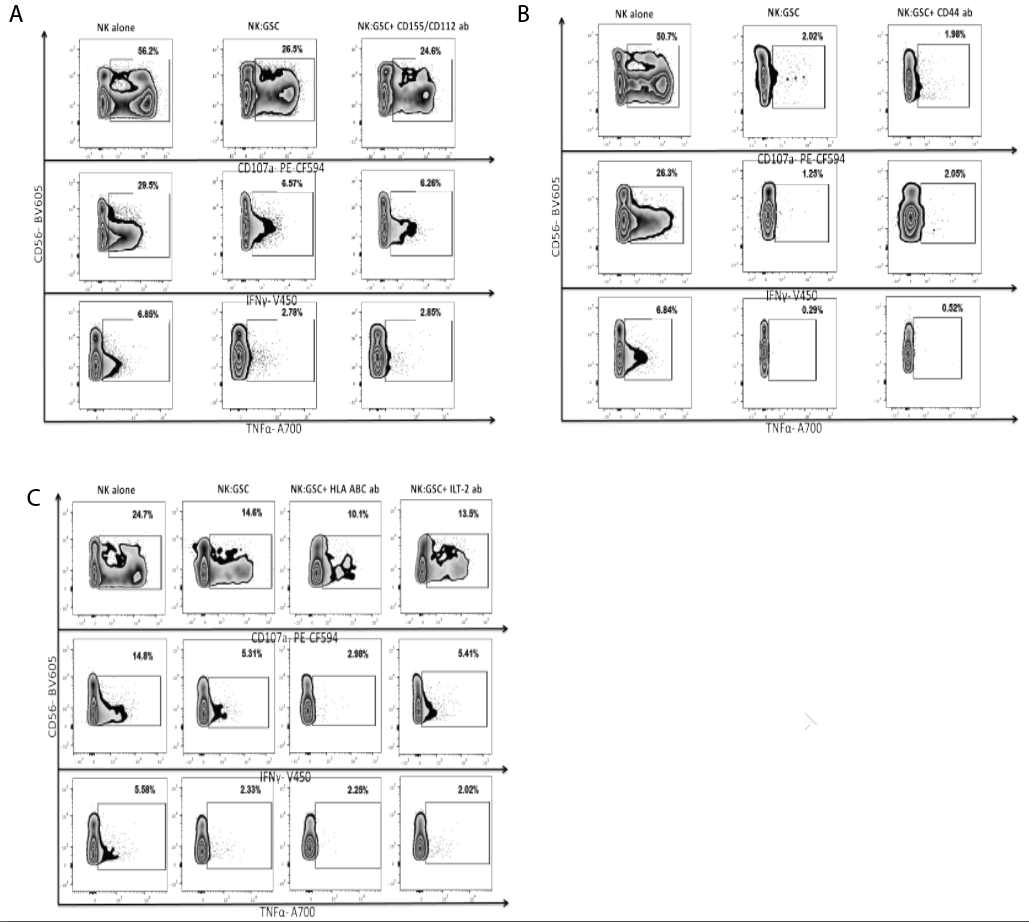

**Supplementary Figure 11. Blocking of major NK cell receptors or their ligands has no impact on GSC-induced NK cell dysfunction. A**-**C**, Representative zebra plots for CD107a, IFN-γ, and TNF-α production by NK cells after culture with or without GSCs for 48 hours in the presence or absence of blocking antibodies against CD155/CD112, CD44, HLA-ABC and ILT-2. NK cells were gated on CD3-CD56+ lymphocytes. Inset numbers are the percentages of CD107a-, IFN-γ- or TNF-α-positive NK cells within the indicated regions.

Supplementary Figure 12

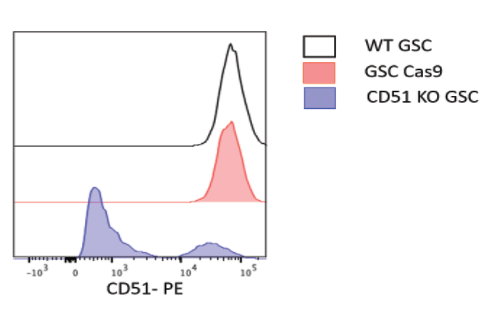

**Supplementary Figure 12. CRISPR/Cas9 silencing of αv integrin (CD51) in GSCs. A,** Representative histograms showing CD51 expression on the surface of wild type (WT) GSCs (white), GSCs treated with CRISPR Cas9 (GSCs Cas9 control; red) or GSCs after *CD51* KO (blue).

Supplementary Figure 13

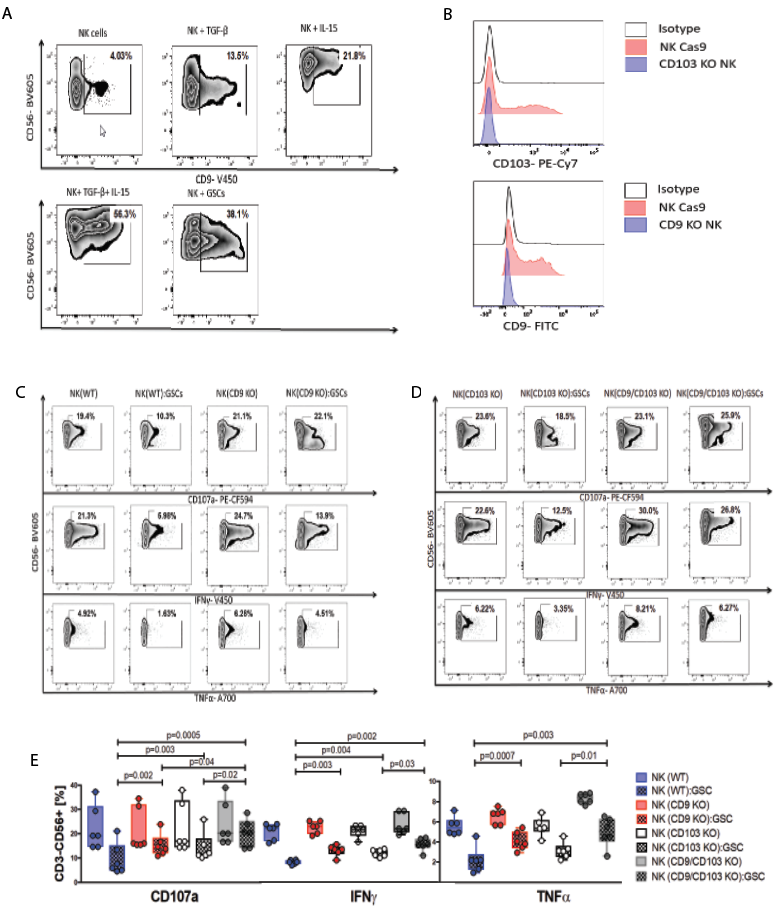

**Supplementary Figure 13. CD9/CD103 expression on NK cells is induced by TGF-β and can be effectively silenced using CRISPR/Cas9 gene editing. A,** NK cells were cultured in SCGM, or in SCGM supplemented with 10 ng/ml TGF-β and/or 10 ng/ml IL-15, or with GSCs in a 1:1 ratio for 48 hours. After 48 hours, the cells were harvested and stained for surface expression of CD9. **B,** Representative histograms showing the expression levels of CD9 (bottom) and CD103 (top) on the surface of NK cells following treatment with CRISPR Cas9 control (red), CRISPR Cas9 *CD9* KO (bottom, blue) or CRISPR Cas9 *CD103* KO (top, blue) as assessed by flow cytometry. **C-E,** Representative zebra and box plots for CD107, IFN-γ, and TNF-α production by WT NK cells, *CD9* KO NK cells, *CD103* KO NK cells and *CD9/CD103* double KO NK cells in response to K562 targets (n=6). Inset numbers are the percentages of CD107a-, IFN-γ- or TNF-α-positive NK cells within the indicated regions. P values were derived using paired t-test.

Supplementary Figure 14

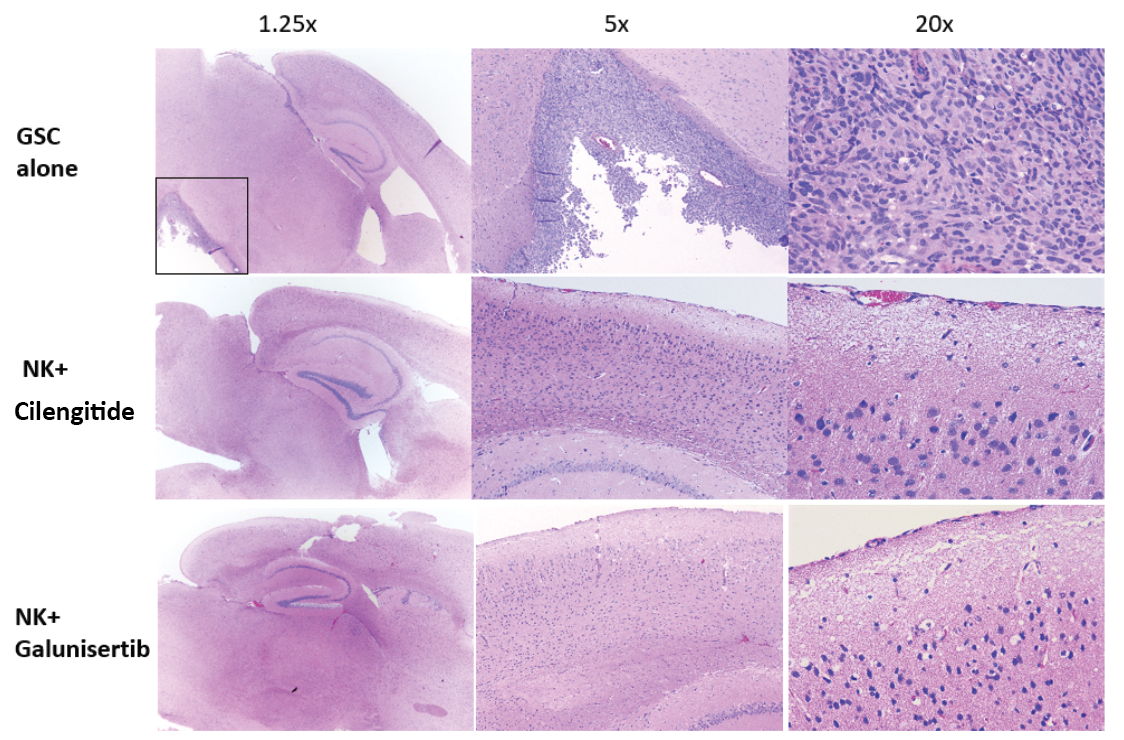

**Supplementary Figure 14. NK cell therapy in combination with galunisertib or cilengitide eliminates glioblastoma in vivo.** Photomicrographs showing severe infiltration and effacement of the cerebral gray matter by glioblastoma in an untreated control mouse in comparison to mice treated with combination therapy with NK cells and cilengitide or galunisertib, which shows no evidence of tumor. (H&E, 1.25x objective; 20x objective inset).

Supplementary Figure 15

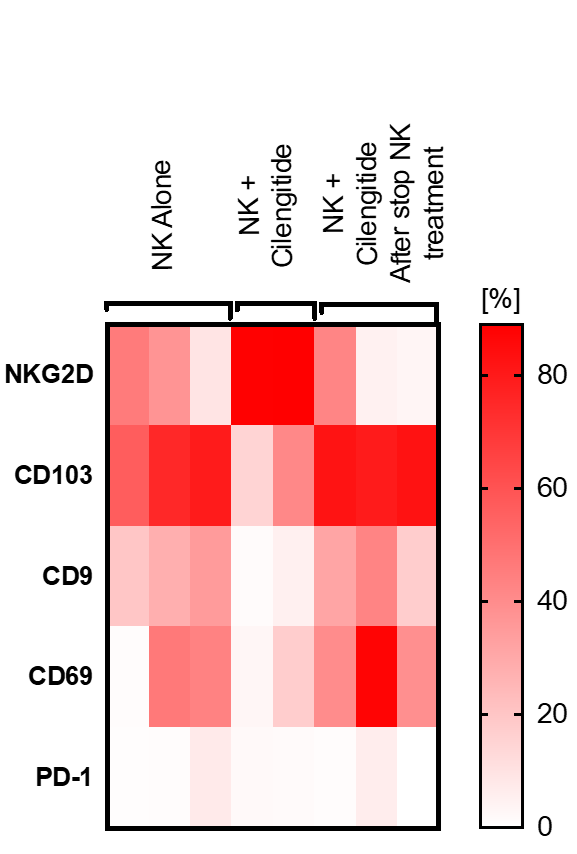

**Supplementary Figure 15. Cilengitide treatment protects NK cells from TGF-β induced inhibitory phenotype in vivo.** Heat map representation of the surface markers NKG2D, CD9, CD103, PD-1 and CD69. Once the mice were sacrificed, the brain tissue was processed and TiNKs were extracted from the GBM tumor. The cells we then stained for the indicated surface markers and analyzed using flow cytometry. The expression is presented as percentage of the cells expressing each marker within the total NK cell population.

Supplementary Figure 16

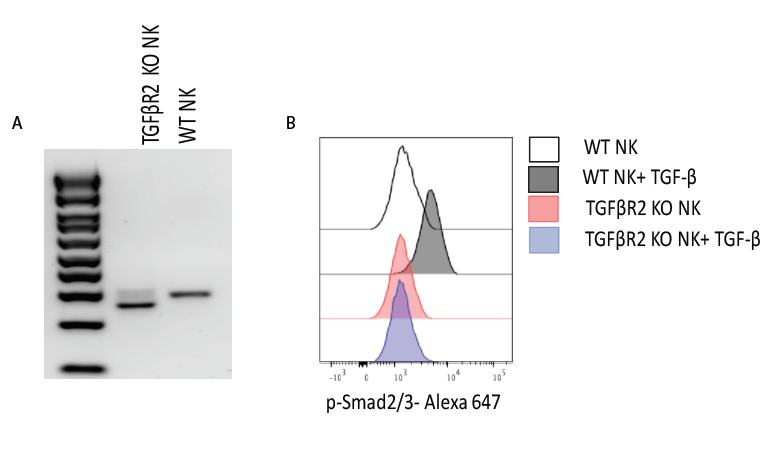

**Supplementary Figure 16. CRISPR/Cas9 silencing of *TGFβR*2 in NK cells. A,** The *TGFβR2* KO efficiency was determined by PCR. **B**, Representative histograms showing abrogation of p-Smad2/3 signaling in *TGFβR2* KO NK cells in response to treatment with exogenous TGF-β (10 ng/ml) for 45 mins compared to WT NK cells.

Supplementary Figure 17

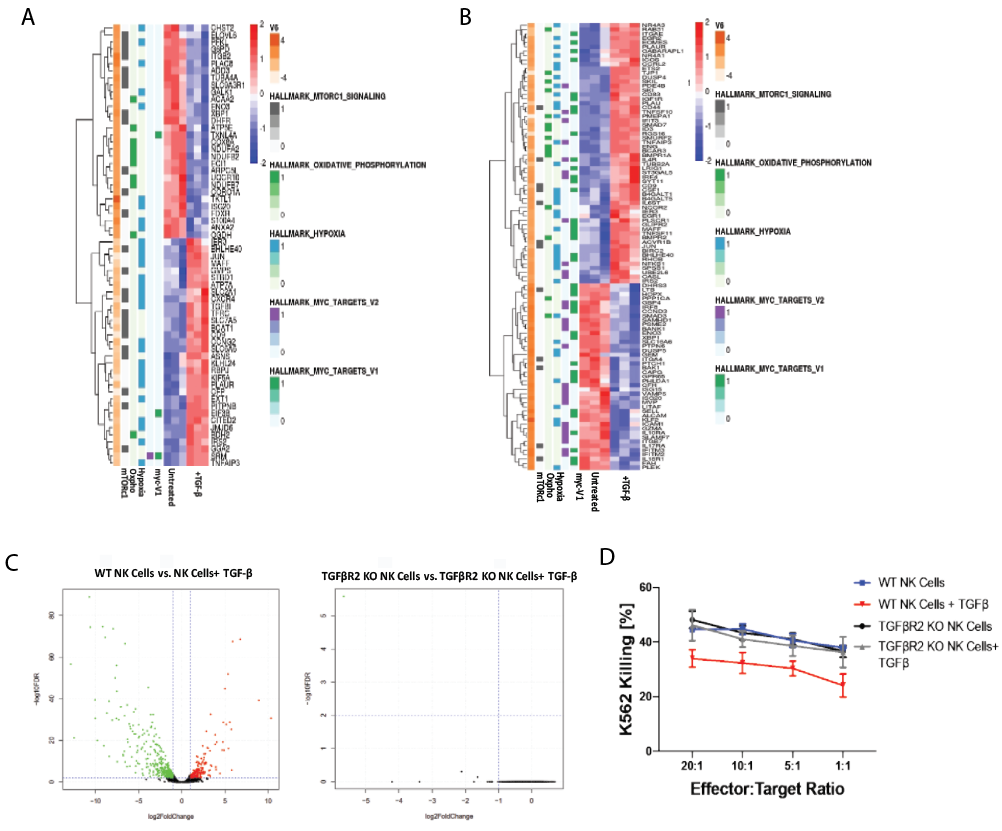

**Supplementary Figure 17. CRISPR/Cas9 silencing of *TGFβR2* in NK cells. A-C,** Transcriptomic analysis of WT-NK and *TGFβR2* KO before and after treatment with exogenous recombinant TGF-β (10 ng/ml) represented by heatmaps and volcano plots. No change in the gene expression profile was noted before and after treatment with recombinant TGF-β in *TGFβR2* KO cells (n=3). **D**, Specific lysis (^51^Cr release assay) of K562 targets by WT-NK (blue), *TGFβR2* KO NK cells (black) or *TGFβR2* NK cells treated with recombinant TGF-β (10 ng/ml) for 48 hours prior to the assay (red and gray, respectively).

Supplementary Figure 18

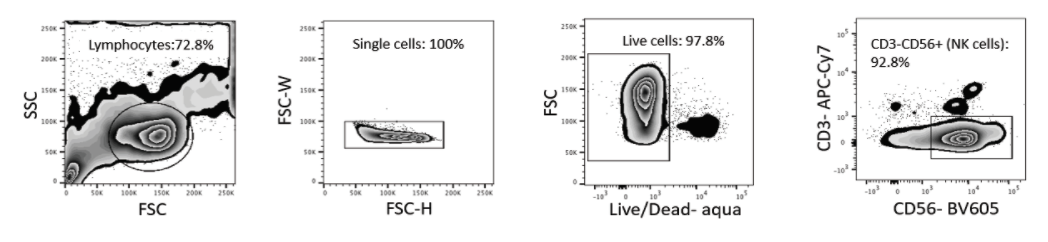

**Supplementary Figure 18. Gating strategy for NK cell phenotyping using flow cytometry.** Representative zebra plots for NK cell gating strategy. Inset numbers are the percentages of lymphocytes, single cells, live cells and NK cells within the indicated regions.

**Supplementary Tables**

Supplementary Table 1. List of antibodies used for mass cytometry.

|  | TARGET | Clone | ISOTOPE | Source |
| --- | --- | --- | --- | --- |
| 1 | CD45 | HI30 | 89Y | Fluidigm |
| 2 | CD57 | HCD57 | 115In | Biolegend |
| 3 | KIR2DL1/S5 | HP-MA4 | 141Pr | Biolegend |
| 4 | EOMES | WD1928 | 142Nd | Thermo Fisher |
| 5 | KIR2DL2/L3 | DX27 | 143Nd | Biolegend |
| 6 | Siglec 7 | EMR8-5 | 144Nd | BD Biosciences |
| 7 | CD62L | DREG-56 | 145Nd | Biolegend |
| 8 | KIR2DL5 | UP-R1 | 146Nd | Miltenyi |
| 9 | CD20 | 2H7 | 147Sm | Fluidigm |
| 10 | TRAIL | RIK-2 | 148Nd | BD Bioscience |
| 11 | SYK | 4D10.2 | 149Sm | Biolegend |
| 12 | KIR2DL4 | 181703 | 150Nd | R&D |
| 13 | CD25 | 2A3 | 151Eu | Miltenyi |
| 14 | CD3Z | 6B10.2 | 152Sm | Biolegend |
| 15 | DAP12 | 406288 | 153Eu | R&D |
| 16 | TIGIT | MBSA43 | 154Sm | Thermo Fisher |
| 17 | CD27 | L128 | 155Gd | Bd Bioscience |
| 18 | KLRG1 | 13F12F2 | 156Gd | Thermo Fisher |
| 19 | CD94 | DX22 | 158Gd | Biolegend |
| 20 | NKP30 | Z5 | 159Tb | Fluidigm |
| 21 | KIR3DL2 | 539304 | 160Gd | R&D |
| 22 | T-BET | 4B10 | 161Dy | Biolegend |
| 23 | NKP46 | BAB281 | 162Dy | Fluidigm |
| 24 | CISH | Polyclonal | 163Dy | R&D |
| 25 | CCR7 | G043H7 | 164Dy | Biolegend |
| 26 | NKG2D | ON72 | 166Er | Beckman Coulter |
| 27 | 2B4 | C1.7 | 167Er | Thermo Fisher |
| 28 | KI67 | Ki67 | 168Er | Biolegend |
| 29 | NKG2A | Z199 | 169Tb | Fluidigm |
| 30 | CD3 | UCHT-1 | 170Er | Biolegend |
| 31 | DNAM | DX11 | 171Yb | BD Bioscience |
| 32 | Perforin | dG9 | 172Yb | Biolegend |
| 33 | Granzyme B | GB11 | 173Yb | BD Bioscience |
| 34 | KIR2DS4 | JJC11.6 | 174Yb | Miltenyi |
| 35 | KIR3DL1 | DX9 | 175Lu | BD Bioscience |
| 36 | CD56 | NCAM16.2 | 176Yb | BD Bioscience |
| 37 | CD16 | 3G8 | 209Bi | Fluidigm |
|  | Cisplatin |  | 198Pt | Fluidigm |

Supplementary Table 2. Sequences used to target *CD9*, *CD103*, *CD51* and *TGFβR2* genes using CRISPR-Cas9 gene editing

|  | Sequence | Exon | Gene Name |
| --- | --- | --- | --- |
| CD9 crRNA | GAATCGGAGCCATAGTCCAA,PAM:TGG | 2 | CD9 |
| CD103 crRNA | GCATTCAAGTGCTGGTCCGG,PAM:CGG | 5 | ITGAE |
| CD51 crRNA | CCACGTCTAGGTTGAAGGCG,PAM:CGG | 1 | ITGAV |
| TGFβRII gRNA | GACGGCTGAGGAGCGGAAGA(gRNA1)  TGTGGAGGTGAGCAATCCCC(gRNA2) | 5 | TGFBR2 |
